# Bion-M 2 Biosatellite: Multisystem Mouse Responses to 30 Days in High-Latitude Orbit as a Deep-Space Analog

**DOI:** 10.64898/2026.05.03.722490

**Authors:** Alexander A. Andreev-Andrievskiy, Mikhail A. Mashkin, Sofya V. Drugova, Vyacheslav A. Shurshakov, Daniil V. Popov, Olga S. Tarasova, Lyudmila B. Buravkova, Olga L. Vinogradova, Vladimir N. Sychev, Oleg I. Orlov, the Bion-M 2 Team

## Abstract

The combined effects of microgravity and deep-space radiation on whole-body physiology remain poorly quantified for future crewed missions. Bion-M 2, a 30-day high-latitude biosatellite carrying group-housed mice, achieved an ISS-comparable total dose with an enriched galactic cosmic ray fraction, approximating conditions beyond low-Earth orbit. A quantitative atlas of 73 physiological endpoints revealed pronounced antigravity muscle atrophy, immune and gastrointestinal remodeling, and delayed recovery of hematologic and visceral indices through 30 days post-landing. A dry-food–hydrogel diet transformed this response into a stress-dominated, densely interconnected physiological state. Pharmacological Nrf2 activation with omaveloxolone preserved hindlimb muscle mass at ground-control levels and protected visceral organs. These findings establish a systems-level baseline for mammalian adaptation to a deep-space-analog orbit and identify diet and Nrf2 activation as tractable countermeasure levers.

## Introduction

Space animal experiments paved the way for human spaceflight. It is the results of these experiments, notably the Bion program, that made long-duration missions in low-Earth orbit possible (*1, 11*). Ambitious plans for human missions to the Moon and Mars may be constrained by unresolved physiological challenges of deep-space conditions (*7, 18, 20*), making renewed animal experimentation essential for developing adequate biomedical support. As of now, the only platform capable of providing such data is the Bion-M program of satellites—or biosatellites—specifically designed for biological research beyond the well-studied Mir and ISS orbits (*1, 12*). These missions offer a uniquely clean view of spaceflight biology: unlike cosmonauts, experimental animals experience the full, unmitigated combination of microgravity and radiation and can be analyzed at organ, cellular and molecular levels, allowing spaceflight-induced changes and countermeasure effects to be disentangled with a precision that is impossible in humans (*9, 23*). Moreover, the biosatellite format enables large-scale, high-throughput experimentation —including dozens of animals and extensive tissue sharing—that is impractical on the ISS, providing statistical power and biological depth beyond crewed platforms. Here we use the Bion-M 2 biosatellite to (i) chart multisystem mouse responses and 30-day recovery after spaceflight on a high-latitude, galactic cosmic rays (GCRs)-enriched orbit, (ii) test how diet reshapes this organism-level program, and (iii) ask whether pharmacological activation of endogenous defense pathways via Nrf2 can remodel spaceflight and readaptation trajectories. Unlike previous Bion-M missions and ISS-based rodent studies, Bion-M 2 was explicitly optimized to approximate deep-space-like radiation composition while enabling systematic, multisystem readouts during readaptation on Earth.

## Results

### High-latitude Bion-M 2 orbit bridges ISS and deep space

The high-latitude orbit of Bion-M 2 (Fig. 1(**A)**) was chosen primarily to shift the radiation composition toward deep-space-like conditions rather than to increase total dose. Beyond Earth’s magnetosphere GCRs) provide most of the radiation dose, whereas in low-Earth orbit trapped-belt particles dominate. By traversing geomagnetic field lines at high latitude, Bion-M 2 enhances the GCR component while reducing trapped protons, thereby better approximating the radiation environment expected for cislunar and interplanetary missions (*14, 18, 27*).

**Fig. 1.**
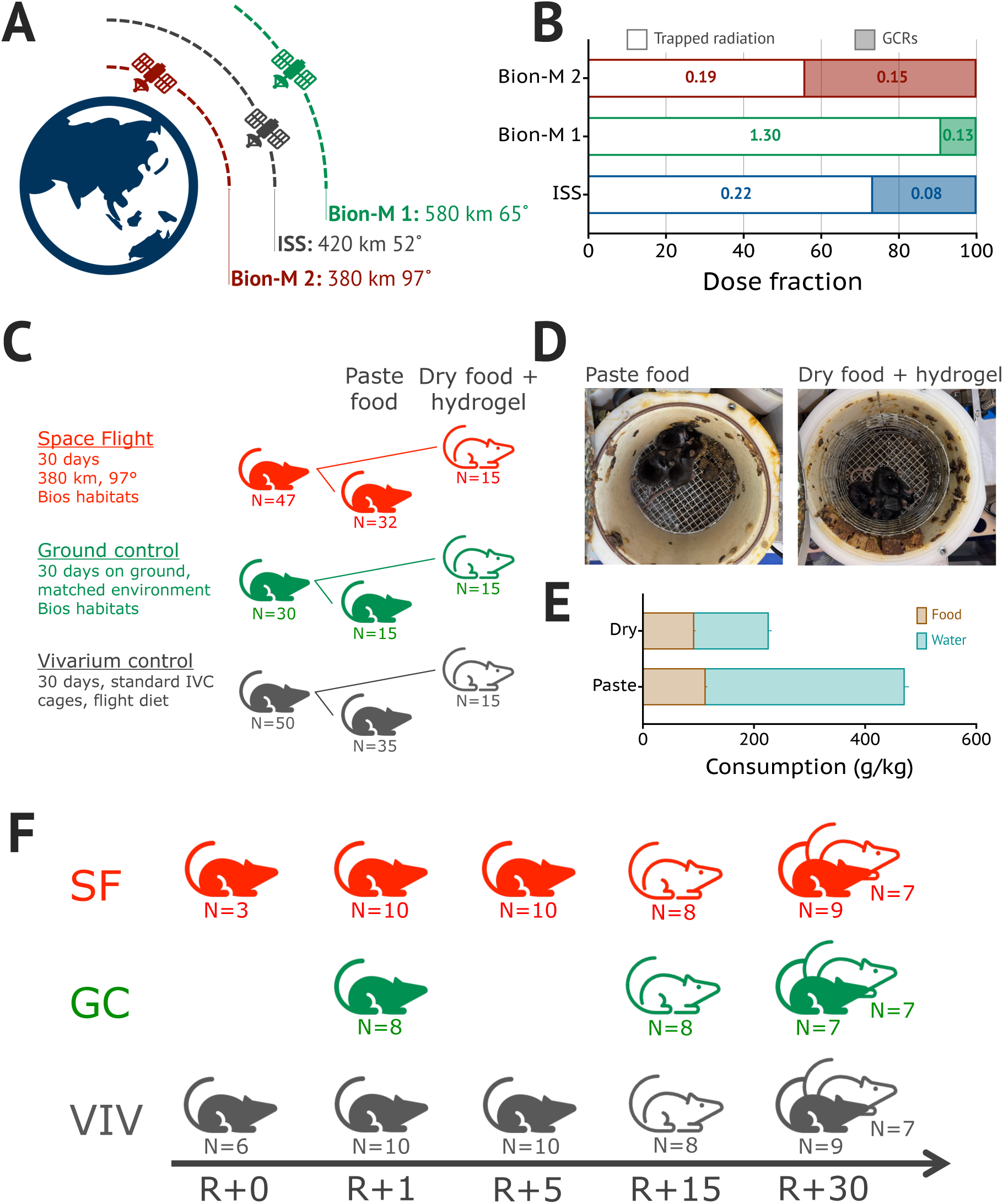
Orbital configuration, experimental design, and post-flight sampling. **(A)** Schematic orbits of Bion-M 2, Bion-M 1, and the ISS with altitudes and inclinations; the high-latitude Bion-M 2 trajectory was chosen to enhance exposure to galactic cosmic rays (GCRs). **(B)** Fraction of total dose produced by trapped protons and GCRs for the three platforms. Stacked bars show contributions from trapped protons (white) and GCRs (shaded); numbers indicate the absorbed dose rate (mGy/day). Bion-M 2 shows an ISS-like total dose but a higher GCR fraction, whereas Bion-M 1 received a larger, proton-dominated trapped-belt dose. **(C)** Experimental groups. Spaceflight (SF), ground-control (GC), and vivarium-control (VIV) cohorts were maintained on paste or dry-food–hydrogel diets in BIOS habitats or standard cages; all animals were individually identified with ear tags and RFID transponders. A parallel Nrf2-modified cohort is shown in Fig. 5A. **(D)** Representative post-flight habitats: dry-food cages appear visibly cleaner than paste-fed ones; mice in both regimens were well-groomed, indicating good overall condition. **(E)** Food (dry weight) consumption was similar for paste and dry diets, but mice obtained 2.7-fold more water from paste than from ad libitum hydrogel. **(F)** Tissue collection schedule, indicating recovery days and animal allocation from each cohort. Paste-fed mice carried implantable temperature loggers, whereas dry-fed mice carried combined temperature and heart-rate loggers for continuous monitoring.

Passive dosimeters inside the Bion-M 2 module recorded an integral absorbed dose of 9 ± 2 mGy, or 0.30 ± 0.06 mGy/day, comparable to the dose measured in the ISS Service Module crew cabin at the current ISS altitude of ≈ 420 km (Fig. 1(**B**)) (*8, 18*). The key difference was composition: relative to ISS, Bion-M 2 showed approximately a twofold increase in the GCR contribution and at least a 30% reduction in trapped-radiation dose (*18*). A moderate X-class solar flare on 26 August 2025 had no measurable impact on onboard dosimetry, underscoring that chronic GCR exposure rather than acute solar events dominated the radiation environment for this mission configuration and duration. Notably, the integral dose on Bion-M 2 was about fivefold lower than during the higher-altitude Bion-M 1 mission (575 km, 64.9°), consistent with reduced trapped-particle flux and minimal exposure to the South Atlantic Anomaly (*1, 8*). Thus, the Bion-M 2 configuration bridges ISS and deep-space radiation environments by achieving a deep-space-like GCR fraction at an overall dose similar to current ISS conditions. We note that, despite its deep-space-like GCR fraction, Bion-M 2 remained shielded by Earth’s magnetosphere; absolute doses and heavy-ion fluences therefore underestimate those expected in true interplanetary transit and should be viewed as a lower-bound analog (*14, 18, 27*).

### Mission overview and experimental design

Within this high-latitude, GCR-enriched orbital environment, Bion-M 2 carried out a 30-day biosatellite mission with mice and other biological payloads, totaling 29 experiments. The satellite was launched on 20 August 2025 from Baikonur and landed on 19 September 2025 near Orenburg after 464 orbits at 370–380 km and 97° inclination. Here we focus on the mouse experiments. Seventy-five mice were housed three per automatic BIOS habitat (25 habitats, spaceflight, SF), with parallel ground-control habitats (GC) and vivarium controls (VIV). This three-tier design — spaceflight in BIOS habitats, ground controls in identical habitats, and vivarium controls in standard cages — disentangles spaceflight per se from hardware- and housing-related effects (*1*).

Male C57BL/6 mice formed the main cohort (Fig. 1(**C**)). In a parallel cohort, we flew Nrf2-knockout mice (*22, 24*) and mice treated with the Nrf2 activator RTA-408 to probe how graded Nrf2 activity modulates spaceflight adaptation (Fig. 5(**A**)). Of the 75 mice launched, 65 survived to landing; 10 animals from the paste-fed C57BL/6J cohort were lost during the final days of the mission due to hardware malfunctions (*1*). Environmental parameters inside the capsule were maintained within the normal range for mice (see Materials and Methods).

The BIOS habitats were newly developed as plastic cylinders with mesh floor and ceiling in two versions: paste-food and dry-food. Paste food (75% water) was delivered at 4-h intervals and served as the sole source of nutrition and water. In the dry-food habitats, food bars lined the walls and hydrogel-based water was delivered into a recessed waterer on the same schedule as paste food. This design generated two contrasting diet and hydration regimens — high-water paste food versus dry food bars plus hydrogel — that could be compared under otherwise identical housing conditions (Fig. 1(**C–E)**).

The main goal of the mouse experiment was to investigate systemic, cellular and molecular responses to prolonged spaceflight and post-flight recovery. To this end, mice underwent a behavioral test battery and automated phenotyping after landing, and tissues from the paste-fed cohorts were collected at landing (R+0) and on days 1, 5 and 30 post-landing (Fig. 1(**F**)). Tissues from dry-fed cohorts were collected after 15 days of behavioral monitoring and again at 30 days post-flight. A tissue-sharing program yielded 158 samples per mouse (> 25,000 samples overall), which were distributed for functional, histological and omics analyses (see Materials and Methods). During the flight, each mouse was continuously monitored with an implantable temperature logger, and cage video was recorded according to a predefined schedule to assess group-housed behavior, generating multi-terabyte telemetry and video datasets for downstream analysis. The resulting telemetry and video datasets extend beyond the scope of this system-level report and will be presented in dedicated follow-up studies.

### System-specific recovery trajectories after paste-fed spaceflight

We first describe the most extensively studied cohort — paste-fed C57BL/6 mice — before turning to integrated analyses and the effects of diet and Nrf2 modulation.

Over 30 days of spaceflight, SF mice gained 7.3 ± 1.6% body mass, closely matching BIOS ground controls (6.1 ± 1.7%) but remaining far below vivarium counterparts (21.4 ± 2.4%; Fig. 2(**A**)). Paste-fed mice thus maintained basic metabolic homeostasis under microgravity yet showed marked growth restraint relative to standard housing. During the 30-day recovery period, SF mice displayed the steepest body-weight gain, significantly outpacing GC and VIV animals (p = 0.031 for Day×Group interaction; detailed slopes in Fig. 2(**B**)), consistent with catch-up growth after the metabolic constraint of spaceflight. A composite normality index derived from short post-flight field videos (mobility, basic sensorimotor responses, external appearance) revealed the expected early deficit in SF mice relative to GC (Fig. 2(**C**)), whereas by the time animals reached the laboratory they had largely regained normal spontaneous mobility and social interaction.

**Fig. 2.**
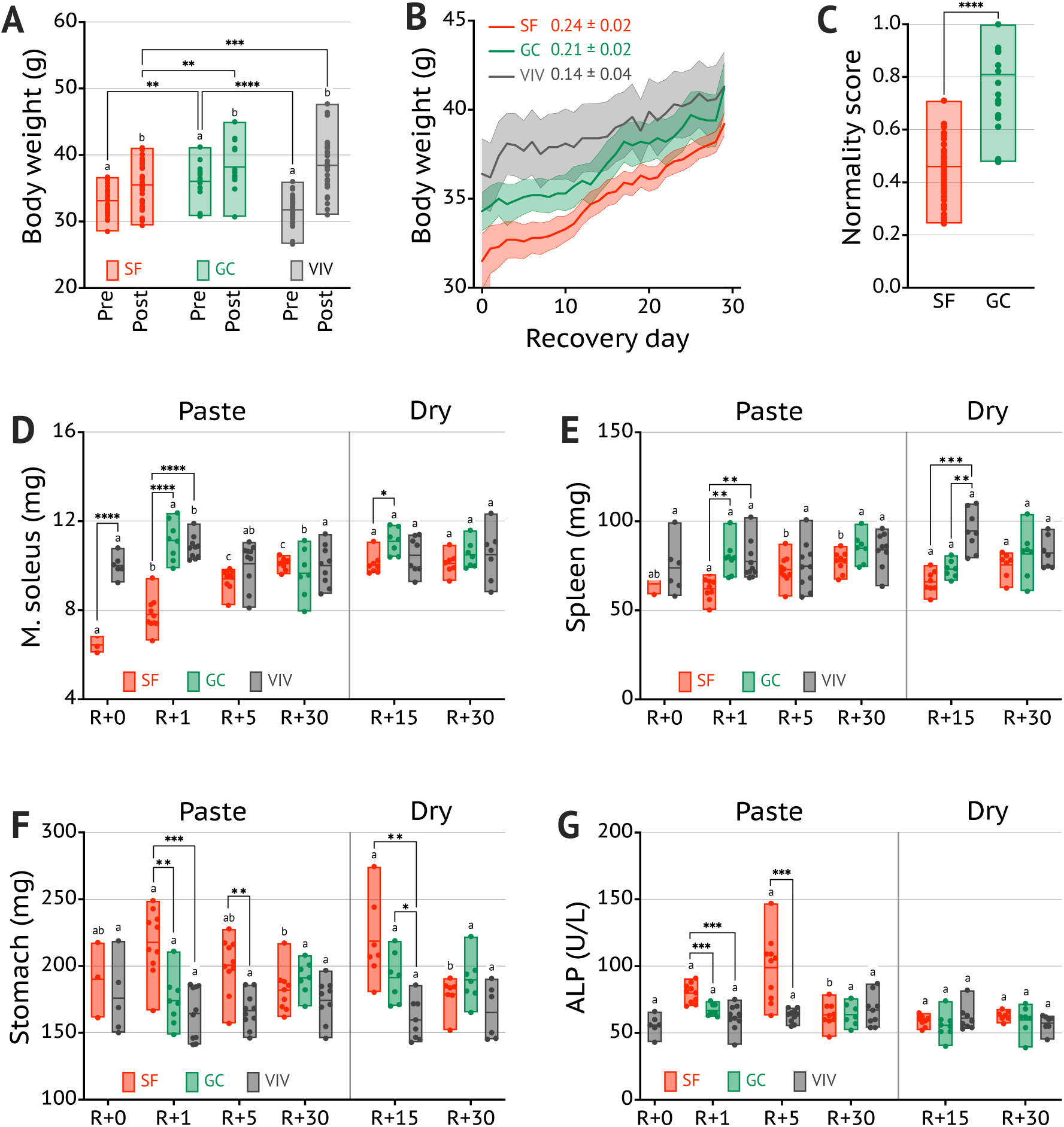
System-specific recovery trajectories after 30-day spaceflight. **(A)** Paste-fed SF mice gained as much weight over 30 days of spaceflight as BIOS ground controls but lagged markedly behind vivarium controls. **(B)** During the 30-day readaptation period, SF mice showed steeper body-weight gain than GC and VIV animals, consistent with catch-up growth. Numbers in the legend indicate weight gain rate (g/day; mean ± sem). **(C)** Composite normality score assessed 3–5 hours after landing, integrating mobility, basic sensorimotor responses and external appearance; SF mice showed significantly lower scores than GC (t = 8.21, p < 0.0001), indicating an early post-flight deficit despite overall good condition. **(D)** Hindlimb muscles showed classic spaceflight-induced deconditioning, with the antigravity m. soleus exhibiting the greatest atrophy and rapid recovery by R+5. **(E)** Spleen mass loss, an indicator of immune suppression, persisted longer after flight than muscle atrophy. **(F)** Stomach mass and other hollow splanchnic organs were markedly increased irrespective of diet, possibly reflecting altered transmural pressure under microgravity conditions. **(G)** Serum alkaline phosphatase (ALP) showed a transient post-flight increase, consistent with early phases of bone and tissue remodeling upon gravitational reloading.

Organ weights revealed the expected pattern of spaceflight-induced deconditioning (*21, 25*). Unless stated otherwise, organ masses were analysed on a log-scale with adjustment for body weight and adiposity. The most pronounced mass loss was observed in m. soleus (−32 ± 2%), the quintessential antigravity muscle, whereas other hindlimb muscles lost 15–20% (Fig. 2(**D**) and Data S1). Spleen mass was reduced by 30 ± 2% (Fig. 2(**E**)), similar to thymus, consistent with spaceflight-associated immune suppression. Stomach mass increased by 27 ± 5% (Fig. 2(**F**)) and colon showed a milder enlargement, indicating gastrointestinal remodeling in microgravity.

The most striking finding at the whole-animal level was that different organ systems recovered on principally different timescales. Most skeletal muscles approached control values by day 5 post-flight, whereas immune and gastrointestinal organs recovered more slowly and often incompletely within the 30-day window (*5, 10, 17*). Hemoglobin and related erythroid parameters were elevated immediately after landing and remained high at least until day 5, compatible with sustained hemoconcentration or altered red-cell homeostasis during early readaptation (Data S1) (*5*). Among blood biochemistry markers, serum alkaline phosphatase was transiently increased in SF mice during the first days of recovery, potentially reflecting enhanced bone turnover and post-flight tissue remodeling (Fig. 2(**G**)) (*10*), and by day 30 most serum indicators had converged with GC and VIV values. To our knowledge, this is the first systematic characterization of post-flight recovery in group-housed mice from landing through day 30, resolving distinct phases of early compensation and later normalization. This divergence between rapid musculoskeletal recovery and delayed immune–gastrointestinal normalization suggests that post-landing performance may be constrained less by muscle strength per se than by slower-recovering stress and barrier systems.

### Integrated multisystem response to 30 days of spaceflight

The organ-by-organ analysis above revealed that spaceflight affects virtually every system we measured, yet some organs change together while others follow independent trajectories. We were particularly interested in the integrated picture — whether spaceflight elicits a coordinated organism-level program or a combination of unrelated organ-specific effects. To address this question, we needed a common currency that would allow us to compare readouts as diverse as muscle mass, leukocyte counts and serum proteins on the same scale. We therefore expressed every endpoint as a standardized effect size (Cohen’s d) — a dimensionless measure of how far a group mean has shifted relative to its variability (*2, 3*). For each of the 73 endpoints, we computed three such effect sizes, each capturing a distinct biological question:

- how much does spaceflight change a given readout compared with vivarium controls living under standard conditions (d_SF–VIV_)?
- how much of that change is attributable to spaceflight per se, beyond the effects of the BIOS habitat (d_SF–GC_)?
- how rapidly does the readout normalize after landing (d_SF–time_)?

Plotting each endpoint in this three-dimensional effect-size space gives a conceptual map in which organs with similar responses to spaceflight and recovery occupy neighbouring regions. For visualization, we projected this three-axis space into two dimensions using UMAP; in this representation, nearby points correspond to endpoints with similar effect-size profiles (Fig. 3(**A**); clustering metrics are shown in in Tables S3, S4).

**Fig. 3.**
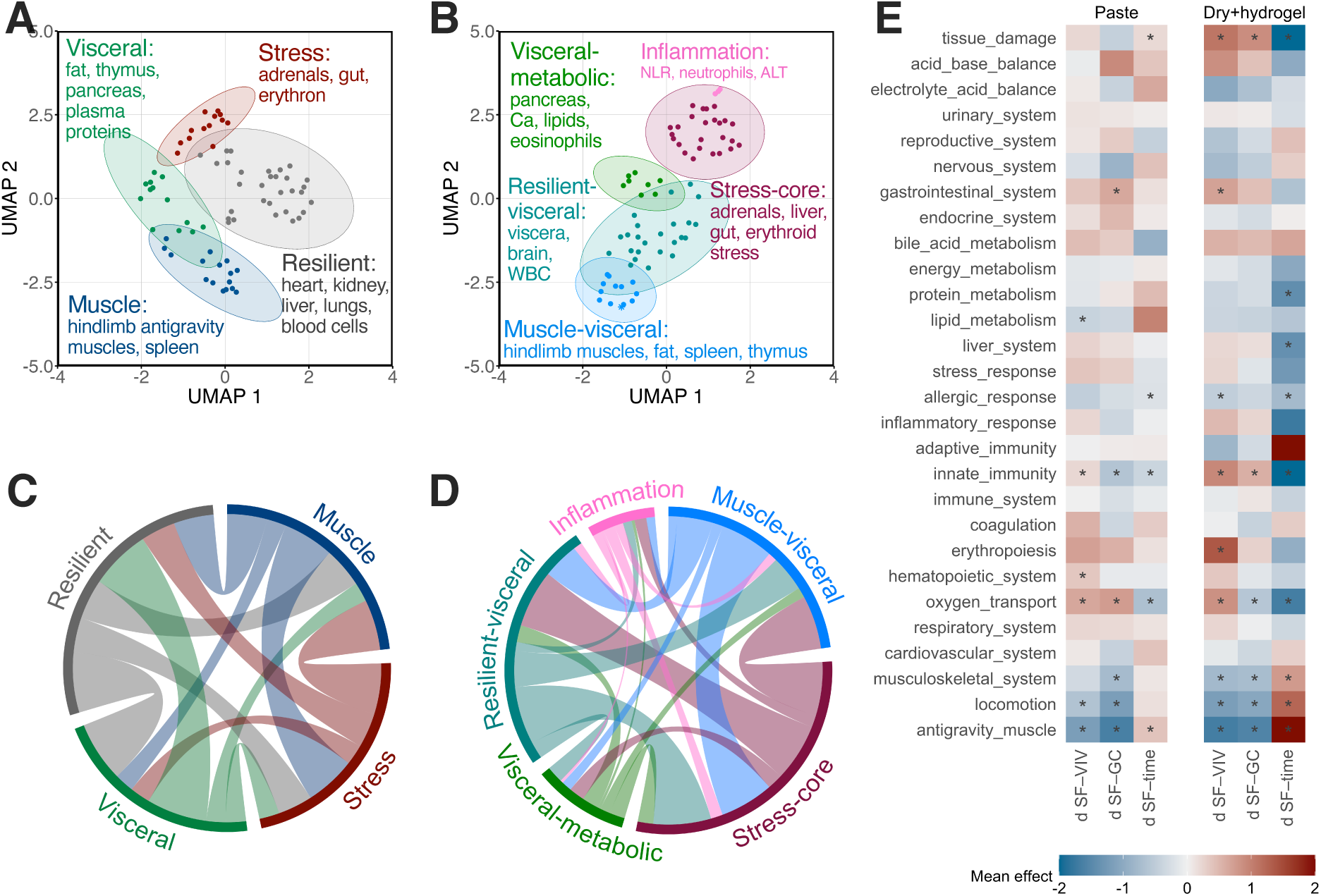
Spaceflight induces a modular physiological response in paste-fed mice that is intensified by a dry-diet challenge. (A, **B)** UMAP plots of 73 physiological endpoints in mice fed a paste (A) or dry, hydrogel-supplemented (B) diet. Points are colored by cluster. The paste diet yields four robust clusters: Muscle (antigravity muscles, spleen, metabolic markers), Stress (adrenals, gut, erythroid indices), Visceral (adipose tissue, thymus, pancreas, serum proteins), and a large Resilient pool. The dry diet partitions the same endpoints into five clusters, adding distinct Inflammation and Visceral-metabolic modules and reshaping the Stress and Muscle-visceral modules, indicating a more complex response. For complete cluster composition and metrics see Tables S3 and S4. **(C, D)** Chord diagrams of inter-cluster connections (|r| ≥ 0.9). Each sector represents a cluster; ribbon width is proportional to the number of edges between clusters. The dry-diet network (D) is strikingly more interconnected, confirming its near-zero modularity (Q ≈ 0) and hyper-integrated architecture. **(E)** Heatmap of mean effect sizes (d) for three orthogonal flight contrasts across 25 functional terms, shown separately for both diets. Terms are systematically grouped; asterisks mark p < 0.05. The heatmap reinforces module-specific signatures: negative musculoskeletal effects, positive immune and stress-related effects, and divergent metabolic profiles. Together, these panels show how a secondary dietary stressor amplifies and reorganizes the spaceflight response, transforming a modular architecture into a densely integrated network.

Four endpoint groups (clusters) emerged consistently across resampling (Fig. 3A). The first, a Muscle module (16 endpoints), links all gravitationally loaded hindlimb muscles with spleen and several metabolic markers. These are the readouts most dramatically reduced by flight and the quickest to rebound on Earth, capturing the canonical deconditioning–reconditioning axis of spaceflight (*10, 21, 25*). A stress module (13 endpoints) brings together adrenal, stomach, colon and erythroid indices — organs that increased in flight and returned to baseline slowly, consistent with a coordinated neuroendocrine, gastrointestinal and hematologic stress program (*5, 17*). A visceral module (13 endpoints) groups adipose tissue, thymus, pancreas and multiple serum proteins (albumin, globulins, lipids), representing a broader visceral compartment with pronounced flight-induced shifts and vigorous post-flight rebound. The remaining endpoints form a large resilient module (31 endpoints) — a relatively stable pool of organs and blood parameters whose net flight effects and recovery slopes are small, indicating robust buffering against 30 days of spaceflight.

When we asked how these four modules relate to each other, correlation-based network analysis revealed that they are not independent compartments but densely connected components of a common physiological response (Fig. 3(**C**); Table S5). The tightest internal coupling was found within the muscle and stress modules (median correlation > 0.9), meaning that if one antigravity muscle atrophied, so did every other, and the same was true for stress-related organs.

We next asked whether these statistically defined modules correspond to recognizable biological programs. Functional annotation confirmed that the muscle module is dominated by antigravity and locomotor functions — not a few idiosyncratic muscles but a coordinated program of gravitationally loaded tissues (*4*). The stress and visceral modules are enriched for innate-immunity genes, consistent with a coordinated neutrophil- and monocyte-driven activation (*5*). Oxygen-transport and hematopoietic annotations captured a transient erythroid elevation during flight followed by post-flight normalization. Terms related to tissue damage exhibited significant time-dependent changes, reflecting an active tissue-repair component of the recovery process.

Together, these analyses delineate a coherent spaceflight-and-recovery program dominated by antigravity muscle wasting, innate-immune activation and organ-system remodeling, rather than a collection of unrelated organ-specific effects.

### Diet shapes organism-level responses to spaceflight

All Bion missions to date have relied on a semi-liquid paste diet. Although proven reliable, the paste has drawbacks: its high-water content (≈ 75%) shifts water–electrolyte balance, increases waste-processing demands, and complicates comparisons with missions using dry chow (*1*). Bion-M 2 therefore introduced a dry-food bar combined with hydrogel-based water delivery — a hydrogel that adheres to the waterer and resists floating in microgravity, reducing contamination risk. Because the two diets differed sharply in water content and digestibility (Fig. 1(E)), they created a natural experiment to test how dietary factors shape whole-body physiology during spaceflight.

Because paste is more digestible, paste-fed mice consistently weighed more than dry-fed animals across all housing conditions (SF, GC and VIV). Importantly, among dry-fed cohorts body weights did not differ between SF, GC and VIV mice (Data S1), confirming that the dry-food–hydrogel regimen provided adequate nutritional support in orbit. The lower body mass on dry diet was largely attributable to reduced fat stores (Fig. 4(**A**)), which we accounted for statistically by adjusting organ weights for both body mass and adiposity.

**Fig. 4.**
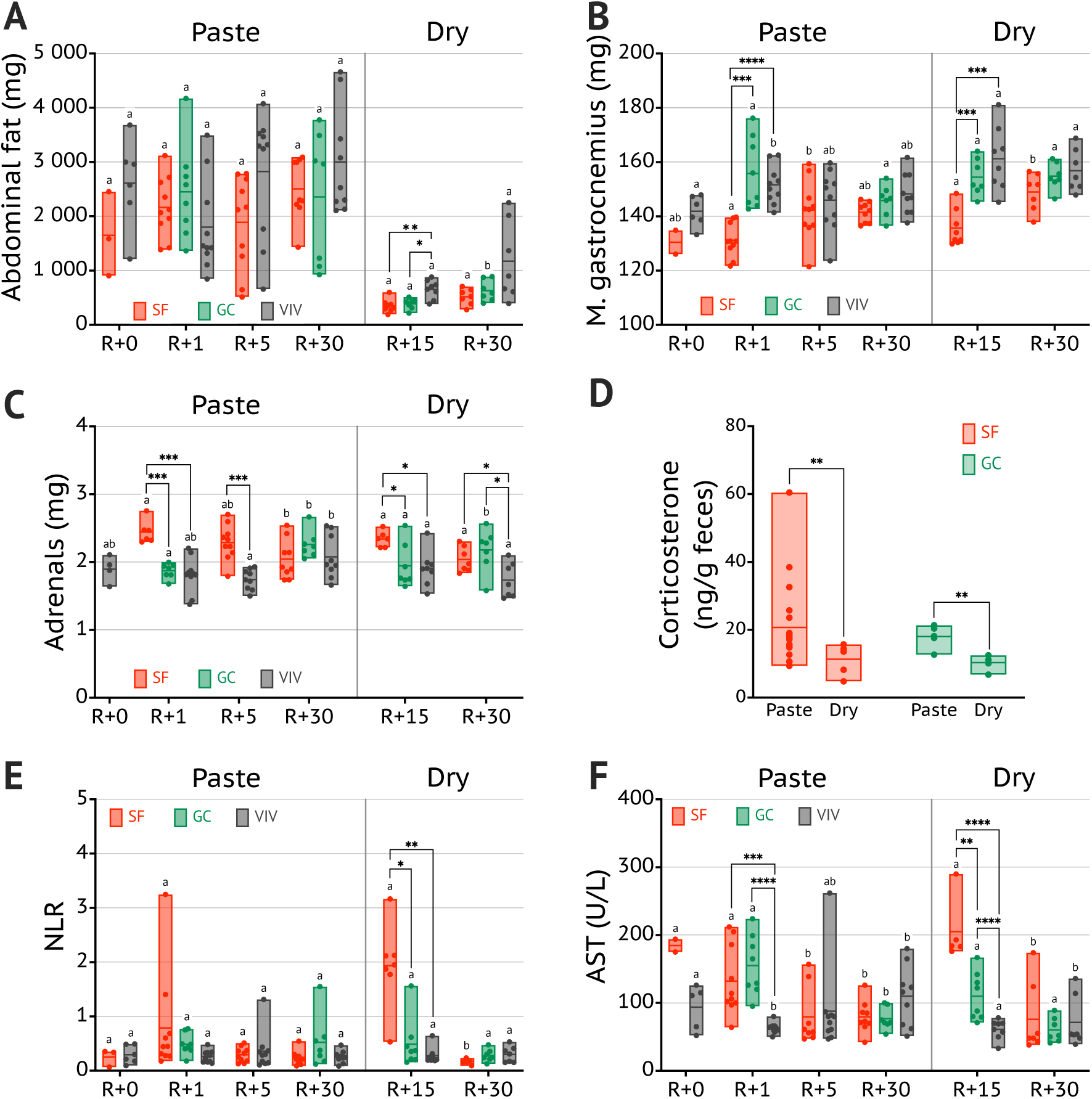
Dry diet modulates spaceflight effects and recovery dynamics. **(A)** Abdominal fat mass in paste- and dry-fed mice across recovery. Lower body weight on dry diet was largely attributable to reduced adiposity. **(B)** M. gastrocnemius mass recovered in paste-fed SF mice by day 5 post-flight but remained reduced at R+15 in the dry-fed cohort. **(C)** Adrenal mass, increased in SF mice, returned toward control levels in paste-fed animals but remained elevated at least through R+15 in the dry-fed group. **(D)** Cumulative fecal corticosterone over the 30-day mission was lower in dry-fed than in paste-fed BIOS habitats under both SF and GC conditions, suggesting the dry-food–hydrogel regimen was well tolerated in orbit. **(E)** Neutrophil-to-lymphocyte ratio (NLR), an index of immunological stress, was elevated in dry-fed mice and normalized only by R+30. **(F)** AST, a marker of liver damage, showed a transient post-flight elevation in paste-fed SF mice but remained markedly increased in the dry-fed cohort as late as R+15, with a graded rise from VIV through GC to SF indicating additive stress from housing and spaceflight. Together, these data show that while the dry-food–hydrogel regimen is tolerated during flight, it prolongs the physiological stress of readaptation after landing, delaying normalization of muscle, endocrine, immune and tissue-damage markers.

Where the two diets diverged was in recovery dynamics. In dry-fed SF mice, muscle loss was still evident at R+15, whereas paste-fed animals had already partially recovered by that time (Fig. 2D, 4(**B**)). Adrenal hypertrophy — a hallmark of chronic stress — persisted longer in SF mice on dry diet (Fig. 4(**C**)), and gastrointestinal remodeling (stomach and colon enlargement) was present in both diets, with diet per se contributing independently of spaceflight (Fig. 2F).

A paradox emerged when we compared stress markers measured during flight and after landing. Cumulative fecal corticosterone — an integrated measure of glucocorticoid output over the 30-day mission — was actually lower in dry-fed than in paste-fed habitats (Fig. 4(**D**)), suggesting that mice were not more stressed during the flight itself. Yet after landing, dry-fed mice showed pronounced neutrophilia (Fig. 4(**E**)) and strongly elevated AST (Fig. 4(**F**)), classic markers of physiological stress and tissue strain. This dissociation implies that dry feeding shifted the stress burden from a tonic, in-flight glucocorticoid exposure to a sharper post-landing stress response (*5*), rather than representing a uniformly “milder” or “harsher” experience. Housing and diet thus emerged as major determinants of the post-landing recovery phenotype.

When we mapped the dry-diet cohort into the same effect-size landscape used for paste-fed mice, the results were strikingly different. Instead of four response modules, five emerged, and their membership diverged sharply from the paste pattern (p < 0.001; Fig. 3(**B**); full statistics in Table S3). Two of these modules were unique to dry feeding: a compact stress-inflammation module linking immune markers with muscle endpoints, and a protein-energy module tying stress organs to platelet and lymphocyte dynamics — circuits that were invisible on the paste diet and apparently unmasked by the metabolic challenge of dry feeding. The remaining three modules broadly paralleled the paste architecture but with notable shifts: the muscle-metabolic module was now more tightly coupled to metabolic markers; a new visceral-gut module grouped gastrointestinal organs with eosinophils and serum proteins, reflecting the direct impact of the dry-food–hydrogel regimen on gut function and protein metabolism; and the resilient pool was less clearly separated from antigravity muscles.

Beyond reassigning individual endpoints to different modules, dry feeding fundamentally rewired their coordination. The correlation network under dry diet was far denser than on paste, with almost every module strongly connected to every other (Fig. 3(**D**); Table S5). In biological terms, this means that under the metabolic stress of dry feeding, the body’s responses lost their modularity: a change in one system — say, muscle — became tightly coupled to changes in immune activation, hematology and gut function, forming a single, stress-dominated physiological state rather than a set of semi-independent adaptations.

Functional annotation reinforced this picture. On paste diet, the dominant functional signature was antigravity muscle wasting, with more modest changes in oxygen transport and gastrointestinal function (Fig. 3(**E**)). Dry feeding preserved the core muscle-wasting signal but reshaped everything around it. Innate immunity, mildly suppressed on paste, switched to strong activation on dry diet with a pronounced post-flight overshoot (Data S1). Oxygen-transport and tissue-damage terms, only mildly affected on paste, developed steep recovery dynamics on dry diet, indicating heightened erythroid suppression and active tissue repair. Additional signals in liver function, electrolyte balance and erythropoiesis pointed to broader metabolic engagement. In short, dry feeding did not simply amplify spaceflight effects — it reorganized them, converting a modular set of organ-specific adaptations into a tightly coupled, stress-driven whole-body response.

### Pharmacological Nrf2 activation mitigates muscle loss in flight mice

Having mapped how spaceflight reshapes the body and how diet reconfigures this response, we asked a translational question: can the spaceflight-induced program be therapeutically modulated? We focused on Nrf2, a transcription factor that sits at the hub of cellular antioxidant and cytoprotective defense (*6, 26*), and that prior ISS experiments had implicated in body-mass maintenance and immune protection during spaceflight (*22, 24*). To span the full range of Nrf2 activity, we flew three groups of mice side by side: Nrf2-knockout animals (deficient defense), untreated C57BL/6 controls (normal defense), and mice treated with the Nrf2 activator RTA-408 (omaveloxolone; enhanced defense) (Fig. 5A). Stepwise induction of the Nrf2 target gene *Nqo1* in muscle and liver confirmed that these three groups achieved graded, functionally distinct levels of Nrf2 pathway engagement in vivo (Fig. 5(**B**)).

**Fig. 5.**
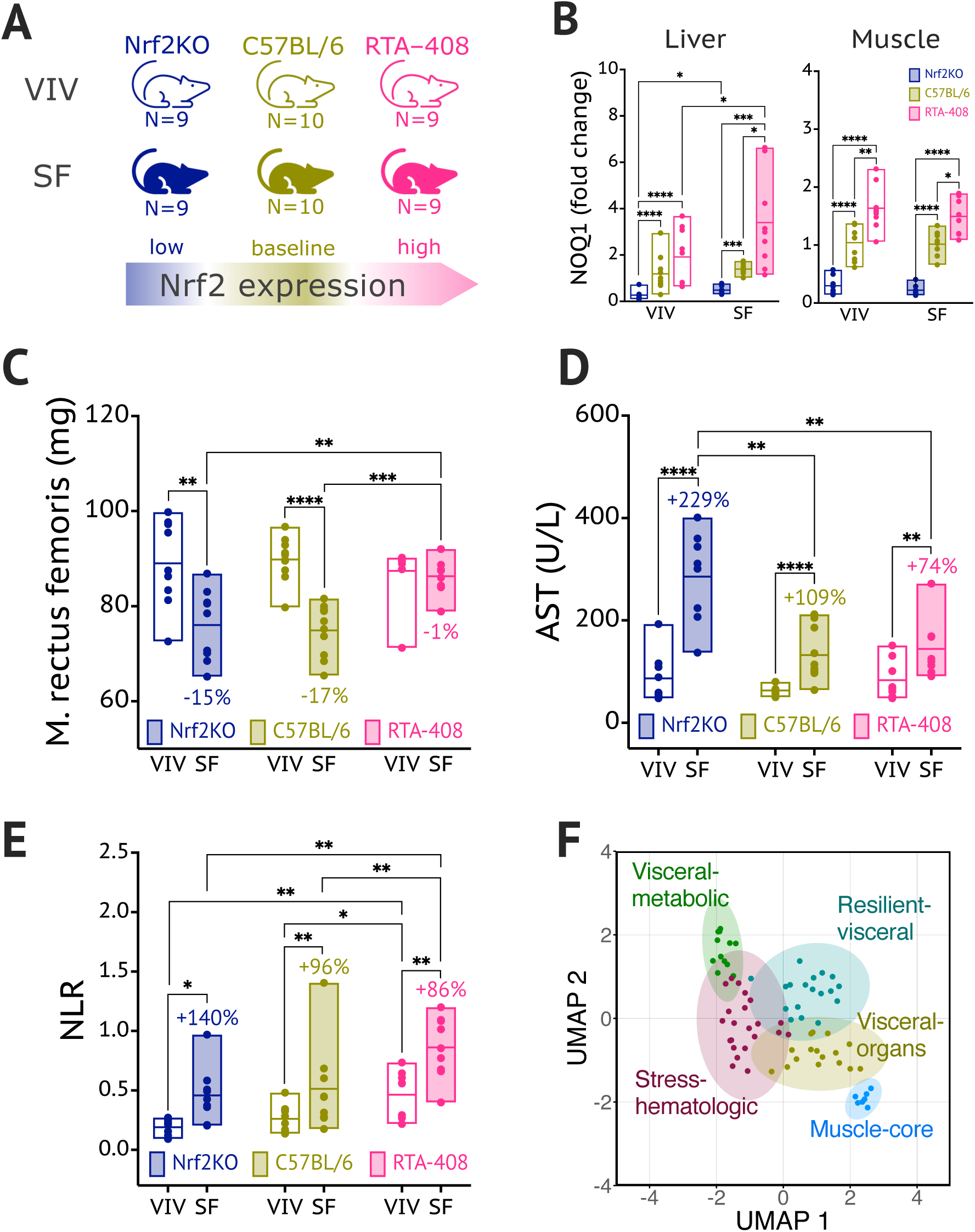
Nrf2 activity level modulates the effects of 30-day spaceflight. **(A)** Schematic design of the Nrf2 experiment: Nrf2-knockout mice (low activity), untreated C57BL/6 (baseline activity), and RTA-408-treated mice (high activity) were flown to span the full range of Nrf2 “dose”. **(B)** *Nqo1* expression in m. rectus femoris increased stepwise with Nrf2 activity under both VIV and SF conditions, confirming effective in vivo titration. **(C)** Body-mass-adjusted m. rectus femoris mass in RTA-408-treated SF mice was indistinguishable from ground controls, whereas untreated C57BL/6 and Nrf2-KO animals showed progressive loss. **(D)** Serum AST, a marker of tissue damage, was negatively related to Nrf2 activity, with the smallest increase in RTA-408-treated mice. **(E)** Neutrophil-to-lymphocyte ratio varied with Nrf2 “dose” in accordance with Nrf2’s role in hematopoiesis, and spaceflight induced similar relative increases in NLR across activity states. **(F)** Clustering of all readouts in the three-dimensional effect-size space yielded modules broadly similar to those in untreated C57BL/6 mice (Fig. 3A), indicating that modulating Nrf2 activity tunes, rather than fundamentally rewires, the organism-level response to spaceflight. Together, these data show that increasing Nrf2 activity protects multiple physiological systems — most notably skeletal muscle—while preserving hematopoietic and immune competence.

The most striking result was near-complete prevention of spaceflight-induced muscle loss by RTA-408 (Fig. 5(**C**)). Untreated C57BL/6 mice lost 18 ± 2% of *m. rectus femoris* mass compared with vivarium controls — one of the largest effect sizes in the entire dataset (Cohen’s d = −2.93). Nrf2-knockout animals ended with the lowest absolute muscle mass of any group. By contrast, RTA-408-treated SF mice retained muscle mass indistinguishable from vivarium and ground controls (Fig. 5C and Data S2). Other hindlimb muscles showed a similar dose-dependent pattern, with partial exceptions in *m. soleus* and *m. vastus intermedius* that likely reflect differences in fiber-type composition or mechanical loading history. Protection extended beyond muscle: kidney mass increased in RTA-408-treated SF mice while declining in untreated and Nrf2-KO animals (*6, 26*), and circulating markers of tissue damage (AST, ALP) were lower in treated mice, pointing to broader organ protection (Fig. 5(**D-E**) and Data S2).

To understand whether Nrf2 simply attenuates spaceflight effects or fundamentally rewires the response, we mapped all readouts into the same integrated framework used above, now asking how each endpoint shifts across the three Nrf2 activity states. The resulting five modules (Fig. 5(**F**); Tables S6, S7) broadly recapitulated the architecture seen in untreated paste-fed mice — a reassuring finding, because it means that Nrf2 modulation selectively tunes the existing physiological program rather than creating a new one.

The muscle–core module showed a striking, non-linear relationship with Nrf2 activity. In relative terms, untreated C57BL/6 mice showed the largest flight-induced loss; Nrf2-KO mice lost somewhat less in percentage terms but started from a lower baseline, ending up with the worst absolute muscle mass of any group. RTA-408-treated mice lost the least mass overall (*15, 16*), effectively preserving their antigravity muscles at ground-control levels — the most pronounced protective effect we observed in this study.

A visceral module — kidney, liver, pancreas, spleen, reproductive tissues and associated metabolites — showed moderate spaceflight-induced changes in untreated mice and a more complex pattern across Nrf2 genotypes. Notably, Nrf2-KO mice were partially protected in some visceral tissues, whereas RTA-408 shifted organs onto a consistently more resilient trajectory, suggesting engagement of broader cytoprotective programs (*6, 22, 24*) beyond simple attenuation of the flight effect.

A stress-hematologic module — overlapping the stress-dominated paste cluster in its erythroid and leukocyte indices, stress enzymes and adrenal mass — showed strong flight responses across all Nrf2 activity states. Nrf2-KO mice mounted a slightly attenuated response, whereas RTA-408 largely preserved the C57-like pattern, indicating that Nrf2 deficiency is subtly detrimental in this axis while pharmacological activation is functionally neutral.

A visceral-metabolic module (plasma proteins, liver-derived metabolites, enzymatic activities) showed robust spaceflight-induced rises in untreated mice that were dampened in both Nrf2-KO and RTA-408 groups. This suggests that intact Nrf2 supports the full metabolic activation response to spaceflight, whereas both deficiency and pharmacological enhancement limit it — although RTA-408 does so from a healthier baseline, preventing excessive stress-driven activation rather than impairing adaptation. The remaining resilient-visceral module contained tissues with small net flight effects; here Nrf2 status shifted baseline set-points without altering the magnitude of the spaceflight response, confirming that these homeostatic systems are largely buffered against both spaceflight and Nrf2 manipulation.

Across this landscape, one finding stands out with particular clarity. Pharmacological Nrf2 activation with RTA-408 protects the systems most vulnerable to spaceflight — antigravity muscles and several visceral organs — while leaving hematopoietic and innate immune competence intact. The protection of rectus femoris mass at ground-control levels, in a 30-day deep-space-analog mission, represents a degree of countermeasure efficacy rarely achieved in spaceflight biology and highlights Nrf2 activation as a promising component of future integrated countermeasure strategies.

## Discussion

For five decades, the Bion program has offered a unique window into mammin alian physiology in space, providing experiments that would be impossible or unethical in humans, yet directly relevant to crewed missions (*1, 11*). Building on this legacy, Bion-M 2 extends animal spaceflight into a high-latitude orbit that increases the contribution of galactic cosmic rays relative to typical low-Earth-orbit missions, while keeping the total dose in the range experienced on the ISS (*8, 18*). Thirty days in this GCR-enriched environment elicited the classical suite of spaceflight adaptations — antigravity muscle atrophy, immune dysregulation, and stress-related organ remodeling — closely paralleling changes documented in cosmonauts and in previous 30-day mouse missions (*5, 10, 17, 21, 25*). By following animals from landing through 30 days of readaptation, we move beyond endpoint snapshots and obtain a systems-level view of recovery, showing that skeletal muscles regain mass within days, whereas immune and gastrointestinal tissues recover more slowly and often incompletely (*5, 10, 17*). This dissociation suggests that the primary physiological bottleneck after a long-duration mission may not be the mechanical ability to move, but the slower resolution of systemic stress and visceral support. For future lunar and Martian crews expected to perform complex tasks immediately after landing, accelerating multi-system recovery — not just muscle strength — becomes a central objective for countermeasure development (*20*).

A central message of this study is that diet is not a neutral backdrop for spaceflight experiments but a major determinant of the organism-level response. Comparing a high-water paste diet (*1*) with a dry-food–hydrogel one, we find that nutritional factors — water content, digestibility, and the resulting gut environment — do not simply shift individual readouts up or down. Instead, they reconfigure how organs and systems move together. On paste, the response to spaceflight was modular: a well-defined antigravity-muscle program, a stress module, a visceral–metabolic module, and a largely buffered homeostatic pool. On dry diet, the same endpoints were pulled into a denser, stress-dominated network in which muscle wasting, innate immune activation, hematologic changes and gut remodeling became tightly linked. The paradox of lower in-flight fecal corticosterone but higher post-landing stress markers under dry feeding fits this picture — rather than experiencing a uniformly milder mission, dry-fed mice appear to trade tonic glucocorticoid exposure in orbit for a sharper, more integrated stress burden during readaptation (*5*). A qualitatively similar pattern of muted in-flight endocrine responses followed by heightened post-flight stress markers has been reported in astronauts, suggesting that nutritional and environmental context can shift when and how the body “pays the price” for spaceflight. These interpretations are tempered by the non-identical sampling schedules for paste- and dry-fed cohorts and by differences in behavioral testing load, which complicate direct comparison of early recovery dynamics. Nevertheless, our findings argue that diet composition should be treated as a primary lever in multi-system countermeasure portfolios rather than as a secondary logistical choice.

Pharmacological activation of the Nrf2 pathway adds a second, mechanistically distinct way to reshape this landscape. Prior ISS missions showed that endogenous Nrf2 contributes to body-mass gain and immune protection during spaceflight (*22, 24*); here we extend this concept to a GCR-enriched orbit and to a pharmacological agonist (*6, 26*). Treatment with the Nrf2 activator RTA-408 markedly reduced spaceflight-induced hindlimb muscle loss, preserving rectus femoris mass at ground-control levels, while Nrf2-knockout mice showed exacerbated atrophy and poorer absolute outcomes. This protection was selective: skeletal muscles and several visceral organs benefited, whereas the characteristic hematologic and innate-immune activation pattern of spaceflight remained largely unchanged. In other words, pharmacological Nrf2 activation did not blunt the entire spaceflight program; it spared the most vulnerable tissues while allowing canonical stress and immune responses to proceed. RTA-408 (omaveloxolone) can modulate other pathways, including NF-!B, in some settings (*6*), and many downstream cytoprotective targets of Nrf2 are subject to additional layers of regulation. Our *Nqo1* expression data and module-level analyses, however, support a view in which the three Nrf2 states sampled here — knockout, baseline, and pharmacologically enhanced — represent distinct points along a stress-defense axis, rather than a simple on/off switch. Within this framework, Nrf2 deficiency and over-activation both dampen an overly strong metabolic activation response, but only pharmacological activation does so from a healthier baseline and with preservation of muscle and visceral integrity.

Viewing all these elements together, Bion-M 2 provides the first integrated analysis of mouse responses to a deep-space-analog mission. At the organ level, we see familiar signatures of spaceflight physiology; at the systems level, we uncover a coordinated spaceflight-and-recovery program in which antigravity muscle wasting, innate-immune activation and organ-system remodeling are tightly linked. Diet and Nrf2 modulation then act as orthogonal control knobs on this program: diet largely dictates how tightly the components are wired together, while Nrf2 retunes the vulnerability and resilience of specific modules within the same architecture. This perspective aligns naturally with emerging human-centric resources such as the Space Omics and Medical Atlas, which emphasize cross-organ and cross-omic integration (*19*). The datasets generated here — including extensive tissue sharing for transcriptomic, proteomic and microbiome profiling — will enable cross-species, multi-omic comparisons of spaceflight adaptation and recovery, helping to identify conserved pathways of resilience and failure.

On note, several design features limit direct extrapolation of Bion-M 2 results to human crews. All mice were male and of a single inbred strain, preventing assessment of sex- and genotype-dependent variability that is likely to be important for human risk prediction (*5, 17*). Group housing in BIOS habitats reduces social-isolation stress but introduces dominance hierarchies and shared microclimates not present in individual astronaut experiences. The 30-day mission samples only the early phase of adaptation; longer-duration flights with mixed-sex cohorts will be needed to chart the transition from acute responses to chronic steady-states and to probe cumulative radiation risks (*14, 27*). The hardware malfunction that reduced the size of the paste-fed SF cohort forced us to cancel the planned R+15 sampling and limits our ability to compare late-recovery trajectories across diets at matched time points. Despite these constraints, the present work establishes a quantitative, systems-level baseline for mammalian adaptation to a radiation environment that bridges low-Earth orbit and interplanetary trajectories. It provides a concrete, testable framework for integrating environmental design (orbit, shielding, diet) with targeted molecular countermeasures (such as Nrf2 activation) in support of human exploration beyond low-Earth orbit.

## Acknowledgments

The authors gratefully acknowledge Svetlana Shipova, Vlada Semenova, Irina Sokolova, and Anna Grigoryan for their contributions to animal preparation, post-flight testing and sample handling. We thank Elena Mednikova for preparing the flight diets. We are indebted to Svetlana Poddubko and her laboratory for microbiological support, and in particular to Kirill Shef for his enthusiastic assistance and for keeping the team’s morale high during critical stages of the project. We also acknowledge the contribution of the dissection team to the success of the tissue-collection program (full list provided in the Supplementary Materials).

We acknowledge the IMBP engineering team, led by Evgeniya N. Yarmanova and Vladislav S. Sedletskiy, for their critical contributions to hardware development and to the success of the mission as a whole.

We thank JSC Experimental Instrument-Making Plant (EPM) for the design and manufacture of the BIOS habitats and associated onboard hardware. We are grateful to Rocket and Space Centre “Progress” JSC and to Alexey Nesterov personally, to the Center for Ground-Based Space Infrastructure Facilities Operation (TsENKI) JSC, to the Federal Air Transport Agency (Rosaviatsiya), to the recovery teams and to the many staff whose daily work made this mission possible.

The authors acknowledge the assistance of Perplexity (GPT-5.1) for English language editing and stylistic suggestions during manuscript preparation; all content was critically reviewed and approved by the authors.

## Bion-M2 Team

Oleg V. Belov^1,2^, Olga V. Fadeeva^3^, Elena A. Markina^1^, Vyacheslav A. Shmarov^1^, Pavel E. Soldatov^1^, Galina Yu. Vassilieva^1^

^1^State Scientific Center of the Russian Federation – Institute for Biomedical Problems of the Russian Academy of Sciences; Moscow, Russia.

^2^ Joint Institute for Nuclear Research; Dubna, Russia.

^3^ MSU Institute for Mitoengineering; Moscow, Russia.

## Author contributions

Conceptualization: AAA, OLV, VNS, OIO.

Methodology: AAA, OLV, VNS, DVP, OST, VAS, LBB, EAM, MAM.

Investigation: AAA, MAM, SVD, VASu, DVP, OST, EAM, OVF, LBB, VNS, OVB, VASm, PES.

Formal analysis: AAA, MAM, SVD, VASu.

Data curation: AAA, SVD, MAM.

Funding acquisition: OIO, VNS.

Project administration: OIO, VNS, OLV, AAA, LBB, SVD.

Supervision: OIO, VNS, OLV, AAA.

Resources: OIO, OLV, VNS, AAA, VASu, DVP, PES, EAM, LBB, OST, OVB, VASm, GYV.

Validation: AAA, SVD, MAM.

Visualization: AAA.

Writing – original draft: AAA.

Writing – review & editing: AAA, OLV, VNS, OIO, DVP, OST, LBB, VASu, MAM, SVD.

## Data, code, and materials availability

All data supporting the findings of this study are available in the main text or supplementary materials. Custom code used for the analysis is available from the corresponding author upon reasonable request.

## Supplementary Materials

### Materials and Methods

#### Experimental groups

As described in the main text, mice were allocated to SF, GC and VIV cohorts of paste-fed C57BL/6J mice sampled at R+0/1/5/30; dry-fed C57BL/6J cohorts sampled at R+15/30; and Nrf2KO and RTA-408-treated cohorts sampled at R+1 (flight and controls). Group sizes and sampling time points are summarized in Table S2. Losses in the paste-fed C57BL/6J cohort were due to a hardware malfunction that caused flooding of one habitat with paste food and subsequent depletion of the corresponding food reservoirs before mission completion. In accordance with pre-defined contingency plans, this reduction in sample size led us to cancel the planned R+15 tissue collection for paste-fed SF mice and to prioritize R+0/1/5/30 time points, as early post-landing days (R+1 and especially R+5) were identified a priori as the most informative window for capturing rapid deconditioning–recovery dynamics. In retrospect, this contingency design choice was justified, as most skeletal muscles and several organ-level endpoints in paste-fed SF mice had largely normalized by R+5, whereas later time points mainly reflected slower adjustment of immune and visceral systems.

#### Mission design and spaceflight environment

The Bion-M 2 biosatellite (Roscosmos) was launched on a Soyuz-2-1b rocket from the Baikonur Cosmodrome (site 31/6) on 20 August 2025 at 20:13 local time (17:13 UTC). After a 30-day mission and 464 orbits in near-circular low-Earth orbit at 370–380 km altitude and 96.6° inclination, the descent module landed near Orenburg on 19 September 2025 at 11:15 local time (08:15 UTC).

During launch and re-entry, animals were exposed to the standard Soyuz-class ascent and ballistic-descent profiles, with accelerations remaining within the nominal envelope for crewed missions; during orbital flight, residual accelerations were effectively negligible relative to 1 g.

A dedicated ground control (GC) experiment was conducted from September, 3 to October, 7 2025,with a 14 days lag relative to space experiment. For logistical reasons, 10 BIOS habitats were operated in a climatic chamber that replicated the microclimate and gas composition measured aboard Bion-M 2, including the programmed temperature cycle, humidity, ventilation schedule, and gas-exchange settings. Vivarium controls (VIV) were maintained in standard cages in the same facility.

Environmental parameters during spaceflight, the ground control experiment and vivarium housing are summarized in Table S1 and Fig. S1. Briefly, mean air temperature in the flight module was 25.1 ± 0.3 °C, relative humidity 66.3 ± 2.7%, cabin pressure 763 ± 11 mmHg, oxygen concentration 20.9 ± 1.2%, and CO₂ 807 ± 924 ppm. Ground-control habitats were maintained at closely matched pressure and gas composition, with a slightly higher CO2 content (1928 ± 553 ppm), while vivarium rooms were kept at 24.9 ± 1.9 °C and 49.4 ± 7.5% relative humidity.

#### Radiation dosimetry

The radiation environment inside and outside the Bion-M 2 descent module was characterized using a combination of passive thermoluminescent and solid-state track detectors and semiconductor-based active dosimeters.

Passive detectors. Thermoluminescent detectors (TLDs) and solid-state track detectors (SSTDs) were deployed in two configurations of the “SPD-M” instrument. Four “SPD” assemblies, each containing TLDs, SSTDs and biological samples, were placed at locations with different shielding thickness inside the descent module to sample depth–dose distributions, while 32 “Dososphere” assemblies containing only TLDs and SSTDs were mounted approximately uniformly over the inner surface of the spherical cabin. Together they provided integral measurements of absorbed and equivalent dose, dose rate, radiation quality factor and linear energy transfer (LET) spectra in the range 10–700 keV/μm, with measurement uncertainty not exceeding ±10% after post-flight calibration.

Active detectors. Semiconductor-based dosimeters monitored temporal dynamics and spectral properties of the radiation field. Two “RD3-B3” units (one inside the cabin, one mounted externally in a payload container) measured charged-particle flux (0.01–1250 cm⁻²·s⁻¹), absorbed-dose rate (0.04 μGy·h⁻¹ to 0.18 Gy·h⁻¹) and energy-deposition spectra corresponding to 0.3–37 keV/μm in water, with maximum uncertainty of ±20%. An additional charged-particle spectrometer, “Tritel-B”, installed inside the cabin, consists of three telescopes of two silicon detectors each (thickness ∼300 μm, sensitive area 150 mm²). Tritel-B measures LET spectra from 0.2 to 120 keV/μm, particle flux from 1.5×10⁻¹ to 4.5×10⁴ cm⁻²·s⁻¹, dose rates from 0.04×10⁻⁶ to 1.0 Gy·h⁻¹ and energy deposition from 0.07 to 83 MeV, allowing calculation of absorbed and equivalent dose. All active instruments recorded data continuously throughout the mission to internal non-volatile memory.

The combined passive–active dosimetry suite enabled reconstruction of absorbed and equivalent doses inside and outside the cabin, LET spectra over a broad range, and separation of galactic-cosmic-ray and trapped-belt contributions based on LET spectra and the directional information from the Dososphere assemblies.

#### Habitats

BIOS habitats were newly developed by SC “EPM” (Saint Petersburg, Russia) specifically for the Bion-M 2 mission in two configurations, “paste” and “dry”, differing only in diet. The habitats provided automated feeding and watering, lighting, waste removal and video recording.

The interior of each habitat consisted of a white caprolon cylinder (diameter 18 cm, height 14 cm; floor area 270 cm²) with stainless-steel mesh (6 × 6 mm) top and bottom. Automated 12 h:12 h light–dark cycles (lights on at 08:00) with white light (4 000 K) provided ≈60 lux at cage level. During the dark phase, infrared LEDs were activated during video recording.

In “paste” habitats, semi-liquid paste food was delivered every 4 h (00:00, 04:00, 08:00, 12:00, 16:00, 20:00) in portions. The default delivery was two 3 g portions per feeding (6 g in total), which could be adjusted from the ground (0–3 portions) to maintain a small excess of food in the cage without overloading the waste-removal system. In “dry” habitats, compressed food bars were lined along the interior wall and supported by a metal mesh (2 × 2 cm). Hydrogel (1.5% medium-viscosity carboxymethylcellulose; Sigma, USA) was delivered into a recessed waterer at the same 4-hour intervals, ≈2.2 ml per watering.

Waste removal was achieved by continuous airflow through the cage (≈0.15 m/s). Feces for corticosterone analysis were retrieved from the waste removal system after the flight or ground control experiment, frozen and stored at −20 °С for subsequent analysis.

All other environmental parameters in the habitats were maintained by the spacecraft or climatic-chamber life-support systems as described above.

#### Home-cage behavior in flight

Home-cage behavior was recorded continuously for the first 7 days of the flight, weekly thereafter (flight days 14, 21 and 28), and during the terminal phase (flight day 30, descent and ≈12 h post-landing). Between these continuous recording sessions, 1 h videos were acquired around each feeding event (15 min before and 45 min after each feeding). Videos of the 20:00 feeding (corresponding to peak activity at “sunset” in the programmed light–dark cycle) were downlinked on average every other day for real-time monitoring of animals’ condition. Across the flight and ground-control experiments, a total of 9,460 h (33.1 TB) of video was collected for subsequent analysis.

#### Animals

Mice (all males) were flown at 5 months of age. Two mouse strains were used: C57BL/6 and Nrf2-knockout (Nrf2KO). All animals were specific pathogen-free (SPF). C57BL/6 mice were obtained from the Center for Laboratory Animals Genetic Resources (Institute of Cytology and Genetics, Russian Academy of Sciences, Novosibirsk, Russia). A homozygous Nrf2KO stock (strain C57BL/6Cya-Nfe2l2em1/Cya) on a C57BL/6 background was purchased from Cyagen (Suzhou, China) and subsequently bred in house at the Institute for Biomedical Problems (IMBP). Nrf2KO mice used for the flight and corresponding controls were genotyped according to the protocol provided by the strain developer.

The study was approved by the Commission for Bioethics of the Institute for Biomedical Problems, Russian Academy of Sciences (protocol № 689, 04.08.2025), and complied with national and guidelines for the care and use of laboratory animals.

Animal training for spaceflight was performed as described for the Bion-M 1 mission [1]. Briefly, all mice were individually marked with ear tags (RWD, China) and subcutaneous RFID transponders (8 × 1.4 mm, ICBC, Zelenograd, Russia). Mice were co-housed in trios for at least 2 months before the flight or ground-control experiment, with daily examinations that included body-weight measurements, clinical condition scoring, and assessment of group co-habitation. Trios with stable social hierarchy and without signs of excessive inter-male aggression were selected for further use. Approximately 1 month before the flight, mice were transitioned to the respective flight diets (paste or dry-food–hydrogel). Food and hydrogel consumption were measured daily in the home cage.

During preflight training, mice were housed in individually ventilated cages (500 cm² floor area, Tecniplast, Italy) with wood-chip bedding. To enrich the environment, plastic nesting chambers (Tecniplast, Italy) and shredded paper filler for nesting were provided.

SF mice were transported to the Baikonur Cosmodrome 7 days before launch and housed in open cages in a cleanroom facility. Three days before launch, immediately prior to transfer into the BIOS habitats, the final selection of the flight cohort was performed based on data collected during preflight training and on each trio’s response to transportation and exposure to the novel housing conditions.

#### Surgical procedures and implants

For temperature and heart-rate monitoring, mice were implanted subcutaneously with nanoT or microHRT loggers (Star-Oddi, Iceland) under isoflurane anesthesia (1.5–2% in air) 2 weeks before launch. After preparation of the interscapular field, a subcutaneous pocket was created over the pelvic area (nanoT) or on the side of ribcage (microHRT), the device was inserted and the incision closed with 5-0 sutures and tissue adhesive; mice received meloxicam (2 mg/kg, s.c., twice daily) for 2 days post-surgery.

For sustained delivery of RTA-408, silastic implants containing 15% (w/w) RTA-408 in medical-grade silicone elastomer (Flexilis G730, Italy) were prepared and implanted subcutaneously during the same procedure, into a separate pocket on the contralateral side. The silicone–drug mixture was degassed by centrifugation at 5,000 g for 15 min, cured in 0.5-ml syringes and cut into 25-mm rods (2.5-mm diameter). In vitro release was characterized in a 10% DMSO/PBS sink by UPLC–MS/MS quantification of RTA-408 in the medium, and in vivo target engagement was verified by induction of the Nrf2-responsive marker NQO1 in liver and skeletal muscle.

#### Activities at landing site and transportation

Immediately after landing, mice underwent a brief clinical assessment at the landing site. Animals were weighed, simple somatosensory tests (aerial drop test and forepaw placement) were performed and video-recorded, and 30–45-s videos of spontaneous behavior on a tabletop were acquired and scored off-line as described in Data Analysis. After this assessment, mice were returned to their BIOS habitats and transported by air to the Institute for Biomedical Problems (IMBP). Transport and handling conditions (temperature-controlled cabin, minimized vibration and noise) were kept within the same ranges as during preflight transfers. Upon arrival at IMBP, 15 h after landing, animals were transferred to standard individually ventilated cages (as described for preflight housing) for the duration of the post-flight recovery period. Similar to [1], mice of the dry-food–hydrogel cohort were passed through a test battery on R+1 followed by 14-day monitoring in Phenomaster (TSE Instruments, Germany). The details of this extended behavioral analysis will be reported elsewhere.

#### Tissue collection

Tissue collection was performed according to a comprehensive pre-developed necropsy plan yielding 158 samples per mouse and ≈25,000 samples in total (Fig. S2A). Briefly, mice were fasted for 3–4 h prior to euthanasia (except for the 3 animals sampled at the landing site). After collection of fur samples, buccal epithelial smears and other superficial specimens, mice were anesthetized with isoflurane (5% in air), and blood was obtained by right-ventricle puncture via a transdiaphragmatic approach. Blood was immediately divided into aliquots: EDTA-stabilized blood was analyzed for hematological parameters using Dymind DF50 Vet 5-diff research-grade analyzer (China); heparinized blood was used for clinical chemistry on a SMT-120 analyzer with extended reagent disks (Seamaty, China). The remaining blood was allowed to clot in clot-activator and separator gel microtubes for 15-30 min, serum was separated, aliquoted and archived at −80 °C.

After euthanasia, mice were processed by a necropsy team comprising a gross dissector and dedicated subgroups responsible for the head and brain, peripheral muscles, gastrointestinal tract, liver, reproductive organs, skin and bone, and bone-marrow processing. Samples were routed either to immediate functional assays or to cryoprotection and freezing for long-term storage. The complete necropsy procedure for one mouse lasted, on average, 43 ± 3 min (Fig. S2B). Two teams worked in parallel to speed up the process.

To verify that this workflow preserved nucleic acid quality, we measured RNA integrity numbers (RIN) in hindlimb muscles collected immediately after sacrifice or after a 40-min delay, matching the upper range of our necropsy times; RIN values did not differ between conditions and were consistently around 9.

For each sampling time point, SF, GC and VIV animals (and, where applicable, Nrf2KO and RTA-408 cohorts) were euthanized and processed in parallel. Mice from different cohorts were randomly interleaved across dissection lines and operators, so that each line handled a mixture of groups rather than a single cohort. This design minimized potential batch effects associated with dissection order, team and clock time, and ensured that space-flight and control animals experienced comparable handling conditions throughout tissue collection.

#### Fecal corticosterone

Fecal corticosterone was quantified in samples collected over the 30-day SF and GC experiments from the waste-removal system of BIOS habitats. Fecal pellets were air-dried at room temperature for approximately 24 h and manually cleared of food debris, then additionally dried at 60 °C overnight and ground to a fine powder. For extraction, ≈1000 mg of fecal powder was mixed with 1 ml of 80% methanol, incubated for 1 h on a rotation shaker and centrifuged at 15 000 rcf. The supernatant was mixed 1:1 (v/v) with water, incubated at −20 °C to precipitate contaminants and centrifuged again; clarified extracts were transferred to low-sorption tubes and dried in a vacuum concentrator (Concentrator plus, Eppendorf, Germany). Residues were washed with 80% methanol, dried again, reconstituted in 80% methanol, diluted 1:20 in PBS and assayed using competitive corticosterone ELISA kits (Cloud-Clone, China) according to the manufacturer’s instructions on an A400 microplate reader (AllSheng, China). The extraction and assay procedure was partially validated by spike-in recovery experiments, confirming accurate quantification of exogenous corticosterone added to fecal extracts.

#### Reverse transcription and quantitative PCR

Total RNA was extracted from frozen liver and skeletal muscle using column-based kits (Evrogen, Russia) according to the manufacturer’s protocols. RNA quantity and purity were assessed spectrophotometrically (ND-1000, NanoDrop, USA); only preparations with A260/280 and A260/230 ratios of 2.00 ± 0.05 were used for downstream analysis.

cDNA was synthesized with the MMLV RT kit (Evrogen, Russia) following the manufacturer’s instructions. An equimolar mixture (10 µM each) of oligo(dT) and random hexamer primers (Evrogen, Russia) was used, and 200 ng of total RNA were reverse-transcribed per reaction for 1 h at 37 °C. The resulting cDNA was diluted in TE buffer to 12.5 ng/µl and stored at −80 °C until use. Quantitative PCR was performed on a DTlite cycler (DNA Technology, Russia) using qPCRmix-HS SYBR master mix (Evrogen, Russia); 20 ng cDNA were added per reaction. Expression levels of the target gene Nqo1 (forward 5′-GTCCTCCATCAAGATTCG-3′, reverse 5′-GCTAACTGCTAACTGCTAA-3′) were measured relative to the reference genes Hprt (forward 5′-GCAGTACAGCCCCAAAATGG-3′, reverse 5′-AGTCTGGCCTGTATCCAACAC-3′) and Gapdh (forward 5′-AGGTCGGTGTGAACGGATTTG-3′, reverse 5′-TGTAGACCATGTAGTTGAGGTCA-3′) using the ΔΔCt method.

#### Data analysis

All analyses were performed in R (version 4.3.2). Continuous endpoints were analyzed on the natural-log scale, which provided the best approximation to normality and homoscedasticity. For organ weights, we explicitly accounted for group differences in body size and adiposity: models included body weight and fat mass as covariates, and figures report covariate-adjusted (least-squares) means and confidence intervals on the original scale (obtained by back-transformation from the log scale). This correction is motivated by systematic shifts in body weight and adiposity between groups (see Fig. 2A,B and Fig. 4A), which would otherwise obscure true organ-specific effects.

Observations with absolute standardized residuals ≥ 3 in initial models were treated as outliers and excluded, after which models were refitted and covariate-adjusted values on the original scale were obtained for descriptive statistics and effect-size calculations. Linear models on the log scale were then used to estimate the effects of group, diet and Nrf2 status; p-values were adjusted for multiple testing using the Benjamini–Hochberg procedure. Standardized differences (Cohen’s d) were calculated on back-transformed, covariate-adjusted means, ensuring that effect sizes are expressed for biologically interpretable values (e.g. milligrams) while still benefiting from modelling on the log scale. Where several recovery time points were available, effect sizes were additionally summarized by random-effects meta-analysis (DerSimonian–Laird). Endpoints were subsequently clustered according to their pattern of variance attributable to group, diet/genotype and their interaction using k-means, with cluster stability assessed by bootstrap resampling and functional enrichment evaluated by Fisher’s exact test.

Behavioral field-video scores were analyzed with generalized linear models for individual ordinal items and with ANCOVA for a composite Normality Score derived from weighted, normalized item scores; details of the scoring scheme, robustness analyses and full statistical workflow are provided in Supplementary Methods.

## Supplementary methods

All statistical analyses were performed in R (version 4.3.2) using a unified pipeline for data preprocessing, covariate adjustment, outlier handling, univariate modelling, effect-size estimation, meta-analysis, clustering and functional annotation.

### Preprocessing, transformations and covariates

For each animal, measurements were assembled together with group (SF, GC, VIV), dietary or genetic subgroup (paste or dry diet; C57BL/6J, Nrf2KO, RTA-408) and time after landing. Left and right values for paired organs were averaged so that each organ contributed one value per animal and time point. Total fat mass was obtained by summing recorded fat depots and was analysed either as a separate endpoint or as a covariate. To stabilise variance and approximate normality, all continuous endpoints were analysed on the natural-log scale, which provided the best overall performance across data blocks. For organ weights, models additionally included body weight and fat mass as covariates; for fat depots only body weight was used, and blood/biochemistry were analysed without covariates unless specified.

### Outlier handling and covariate-adjusted values

For each endpoint we first fitted a linear model on the log scale including the relevant covariates. Observations with absolute standardised residuals ≥3 were considered outliers and excluded; the immediate post-flight paste-fed SF group was exempt from this filter. Models were then refitted, and covariate-adjusted values were obtained as model predictions at average covariate values plus residuals, back-transformed to the original units. These adjusted values were used for descriptive statistics and effect-size calculations, whereas all hypothesis tests were carried out on the corresponding log-transformed data.

### Univariate models and effect sizes

For each endpoint we fitted linear models on the log scale. For the main cohorts, models included fixed effects of group (SF, GC, VIV), diet (paste vs dry) and, where applicable, recovery time; for the Nrf2 experiment, models included group (SF vs VIV) and genotype/treatment (C57BL/6J, Nrf2KO, RTA-408), with interaction terms when appropriate. Type II analysis of variance was used to obtain F-statistics and p-values, and partial eta-squared was used as a measure of variance explained. P-values were corrected for multiple testing using the Benjamini–Hochberg false-discovery-rate procedure. Standardized differences (Cohen’s d) were calculated on the covariate-adjusted values in original units for all relevant between-group contrasts (e.g. SF vs VIV, SF vs GC, VIV vs GC, SF vs pooled controls) and for within-group changes between successive time points. Significance of these contrasts was assessed by t-tests on the log-transformed data with FDR correction within endpoint, diet and contrast type. Where multiple recovery time points were available, effect sizes were additionally summarized by random-effects meta-analysis (DerSimonian–Laird) to obtain a single estimate per contrast.

### Clustering and functional annotation of endpoints

To classify endpoints by their pattern of sensitivity to spaceflight, diet and Nrf2 status, we summarized for each endpoint three quantities derived from the linear models: the fraction of variance attributable to group, to diet/genotype and to their interaction. These three values were used as features for k-means clustering. The number of clusters was chosen using a combination of silhouette width and elbow criteria, and stability was assessed by bootstrap resampling. Cluster structure was visualized using UMAP projections and correlation-based networks. Endpoints were mapped to broad biological categories (e.g. antigravity muscles, immune system, gastrointestinal tract, lipid metabolism), and enrichment of these categories in each cluster was tested using Fisher’s exact test with FDR correction.

### Clustering of animals (selected analyses)

For selected subsets (notably the Nrf2 experiment), we also clustered animals based on their multiorgan profiles. Hierarchical clustering of scaled, covariate-adjusted endpoints (Euclidean distance, Ward’s linkage) was used to assess how strongly the overall phenotype reflected the intended groupings. Agreement between clusters and true labels was quantified by purity measures, χ² tests and Cramér’s V, with bootstrap perturbation to evaluate stability.

### Behavioral data analysis (field videos)

Field videos were scored offline on predefined ordinal scales for posture, locomotion, exploratory behavior and appearance. Individual items were analyzed with generalized linear models appropriate for their scale (binomial or ordinal logistic regression) including Group (SF vs GC) and time from landing to testing as predictors; false-discovery-rate correction was applied across items. To summarize overall clinical status, raw item scores were mapped to the interval and combined into a weighted Normality Score (higher values indicating more normal function), with greater weights assigned a priori to gait and gross locomotion. Group differences in the Normality Score were assessed by ANCOVA with Group and time from landing as predictors. Robustness to the choice of weights was evaluated by repeatedly resampling item weights within predefined ranges and re-estimating the Group effect.Type or paste text here. This should be additional explanatory text, such as: extended technical descriptions of results, full details of mathematical models, extended lists of acknowledgments, etc. It should not be additional discussion, analysis, interpretation, or critique.

## Dissection team

Belova S.P., Borzykh A.A., Butova K.A., Druzhinina A.A., Fadeeva O.V., Galkin G.V., Gogichaeva K.K., Gordienko K.V., Kalashnikov V.E., Kochakova A.A., Kolesnikov P.Y., Kulakova A.M., Kulishenko A.A., Kurochkina N.S., Kuzmin V.S., Lednev E.M., Leleko L.P., Lukicheva N.A., Mamadzhanian A.M., Markova A.M., Paramonova I.I., Sharlo K.A., Sidorenko D.A., Vepkhvadze T.F., Vilchinskaya N.A., Yakupova E.I., Yarmak D.O.

---

**Fig. S1.**
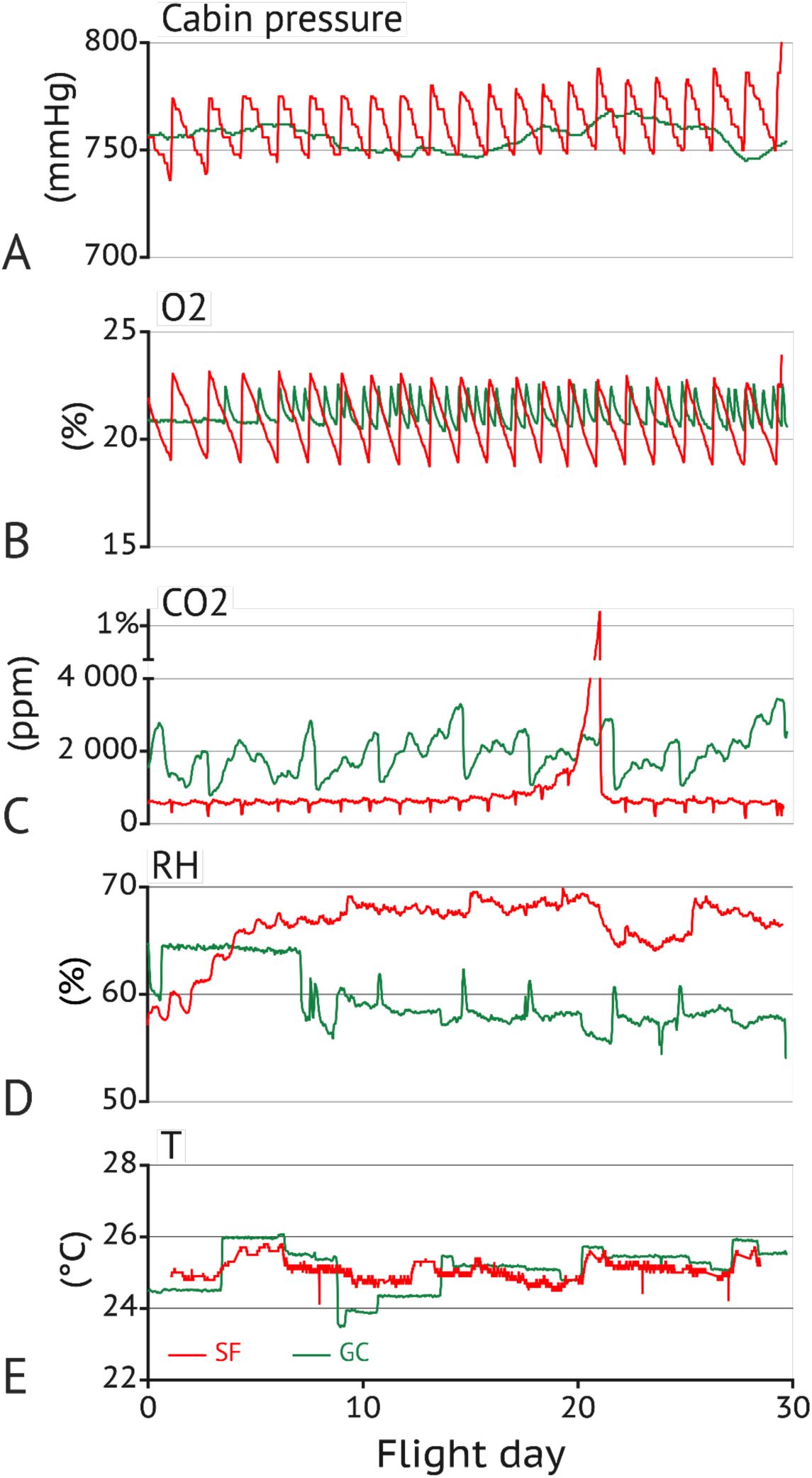
Environmental parameters inside the Bion-M 2 cabin during the 30-day mission. Cabin pressure (A), O₂ concentration (B), CO₂ concentration (C), relative humidity (D) and temperature (E) are shown as 1-h averages over flight days. Note the transient CO₂ elevation up to ≈1% for about a day around flight day 21, which was required to trigger activation of the next CO₂ scrubber cartridge in the life-support system.

**Fig. S2.**
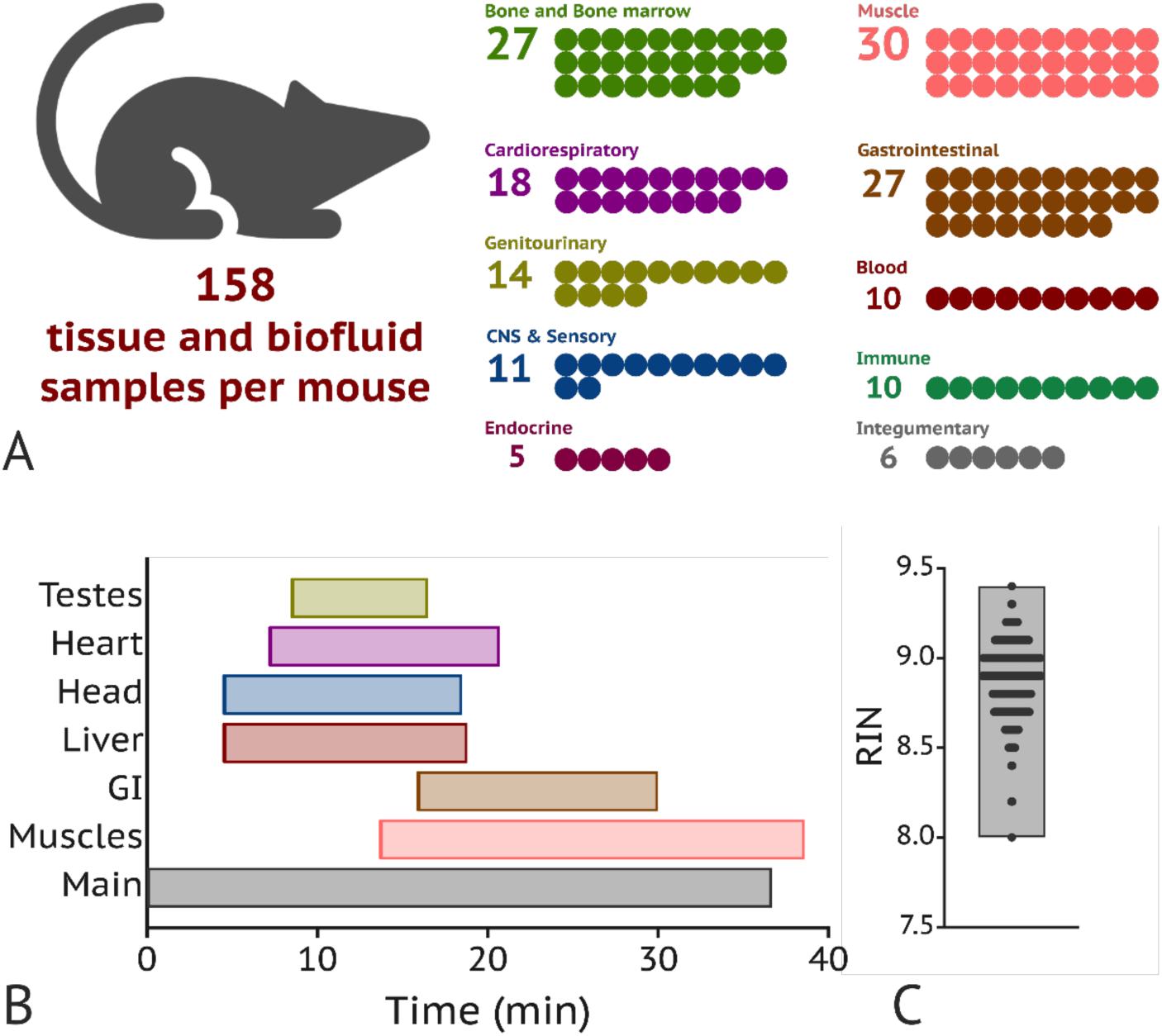
Tissue collection workflow and sample set per mouse. (**A**) Number and distribution of tissue and biofluid samples obtained from each mouse during necropsy, grouped into major organ-system categories. Each dot represents one distinct sample or aliquot. (**B**) Approximate time from euthanasia to completion of the main carcass necropsy (grey bar) and of parallel specialized dissections (colored bars) in paste-fed mice. The main workflow corresponds to whole-body dissection and collection of core organs, while dedicated teams in parallel processed muscle blocks, gastrointestinal tract, liver, head/brain, heart and testes. (**C**) RIN for muscle tissues was 8.9 ± 0.2, indicating good tissue quality for downstream applications despite relatively long dissection.

**Table S1.** Experimental design.

| Group | Diet | Cohort | Size |  | Tissue harvesting |  |  |  |  |
| --- | --- | --- | --- | --- | --- | --- | --- | --- | --- |
|  |  |  | at launch | at landing | R+0 | R+1 | R+5 | R+15 | R+30 |
| SF | Paste | “Paste” | 42 | 32 | 3 | 10 | 10 |  | 9 |
|  |  | Nrf2KO | 9 | 9 |  | 9 |  |  |  |
|  |  | RTA-408 | 9 | 9 |  | 9 |  |  |  |
|  | Dry-hydrogel | “Dry” | 15 | 15 |  |  |  | 8 | 7 |
| GC | Paste | “Paste” | 15 | 15 |  | 8 |  |  | 7 |
|  | Dry-hydrogel | “Dry” | 15 | 15 |  |  |  | 8 | 7 |
| VIV | Paste | “Paste” | 45 | 32 | 6 | 10 | 10 |  | 9 |
|  |  | Nrf2KO | 9 | 9 |  | 9 |  |  |  |
|  |  | RTA-408 | 9 | 9 |  | 9 |  |  |  |
|  | Dry-hydrogel | “Dry” | 15 | 15 |  |  |  | 8 | 7 |

**Table S2.** Environmental and microclimate parameters aboard Bion-M 2 biosatellite, in the ground control experiment and in the animal facility.

| Parameter | Spaceflight | Ground control | Vivarium |
| --- | --- | --- | --- |
| Temperature (°C) | 25.1 ± 0.3<br>(24.2–25.7) | 25.1 ± 0.6<br>(23.9–26.0) | 24.9 ± 1.9<br>(22.2–28.1) |
| Relative humidity (%) | 66.3 ± 2.7<br>(58.3–69.2) | 59.5 ± 2.8<br>(55.9–64.6) | 49.4 ± 7.5<br>(35–61) |
| Atmospheric pressure (mmHg) | 763 ± 11<br>(746–785) | 756 ± 6<br>(746–767) | nd |
| O<sub>2</sub> (%) | 20.9 ± 1.2<br>(18.9–22.9) | 21.2 ± 0.6<br>(20.5–22.5) | nd |
| CO<sub>2</sub> (ppm) | 807 ± 924<br>(317–1455) | 1928 ± 553<br>(997–3181) | nd |

Values are mean ± SD; in parentheses, 2.5th–97.5th percentiles (P2.5–P97.5). nd, not determined.

**Table S3.** Cluster architecture of spaceflight effects in paste-fed and dry-fed mice. For each diet, clusters are numbered as in Fig. 3. Composition lists all physiological endpoints assigned to the cluster. Bootstrap stability (mean ± SD) is the adjusted Rand index (ARI) after 1000 resampling iterations. Mean Cohen’s *d* values are shown for three orthogonal contrasts. At the bottom, metrics comparing the two clustering solutions (paste vs dry) are provided: χ² test, Cramér’s V, agreement rate (%), adjusted Rand index (ARI), and normalized mutual information (NMI). The highly significant χ² (p < 0.001) and low agreement indices confirm that diet fundamentally reorganizes the physiological response to spaceflight.

| Cluster | Composition | Bootstrep<br>stability,<br>m±sd | Mean Cohen's d |  |  |
| --- | --- | --- | --- | --- | --- |
|  |  |  | d<sub>(SF-VIV)</sub> | d<sub>(SF-GC)</sub> | d<sub>(SF-time)</sub> |
| Paste |  |  |  |  |  |
| 1<br>Muscle | Brain, M. rect. fem., M.bulbocav., Spleen, m.gastroc., m.plant., m.quad.fem., m.tib.ant., m.vast.lat., m.vast.med., m. soleus, BUN, BUN/CREA, Ca, LDH, TG | 99±1 | -0.77 | -1.29 | 0.55 |
| 2<br>Stress | Adrenals, Colon, Epididymis, Gaster, Saliv.gl., ALP, Crea, GLU, HCT, HGB, MCH, MCV, RBC | 98±3 | 0.70 | 0.88 | -0.48 |
| 3<br>Visceral | Fat, Pancreas, Thymus, ALB, AMY, GLOB, LPS, TC, TP, tCO2, Lym%, MCHC, PLT | 100±0 | -0.19 | 0.30 | 0.65 |
| 4<br>Resilient | Access.gl., Bladder, Heart, Kidney, LVentr, Liver, Lungs, RVentr, Sept, SmIntestine, Testes, m.vast.intermed., A/G, ALT, AST, CK, PHOS, TB, UA, Eos#, Eos%, Lym#, MPV, Mon#, Mon%, NLR, Neu#, Neu%, PDW, RDW-SD, WBC | 100±0 | 0.23 | -0.29 | -0.22 |
| Dry+hydrogel |  |  |  |  |  |
| 1<br>Resilient-visceral | Bladder, Brain, Epididymis, Heart, Kidney, LVentr, Lungs, M.bulbocav., RVentr, Saliv.gl., Sept, SmIntestine, Testes, m.vast.intermed., m. soleus, ALP, BUN/CREA, GLOB, GLU, TB, UA, Lym#, PDW, WBC | 94±6 | -0.19 | -0.16 | 0.44 |
| 2<br>Inflammation | ALT, NLR, Neu% | 100±0 | 1.29 | 0.75 | -4.04 |
| 3<br>Stress-core | Adrenals, Colon, Gaster, Liver, A/G, AMY, AST, BUN, CK, LDH, LPS, PHOS, TG, tCO2, HCT, HGB, MCH, MCHC, MCV, MPV, Mon#, Mon%, Neu#, RBC, RDW-SD | 98±1 | 0.85 | 0.19 | -1.41 |
| 4<br>Muscle-visceral | Access.gl., Fat, M. rect. fem., Spleen, Thymus, m.gastroc., m.plant., m.quad.fem., m.tib.ant., m.vast.lat., m.vast.med., Lym%, PLT | 99±1 | -1.45 | -0.91 | 1.82 |
| 5<br>Visceral-metabolic | Pancreas, ALB, Ca, Crea, TC, TP, Eos#, Eos% | 98±1 | -0.85 | -0.65 | -1.24 |

$\chi^2$ (12) = 35.43, p = 0.0004; Cramér's V = 0.402; Agreement rate = 12.3%; ARI = 0.096; NMI = 0.184

**Table S4.**
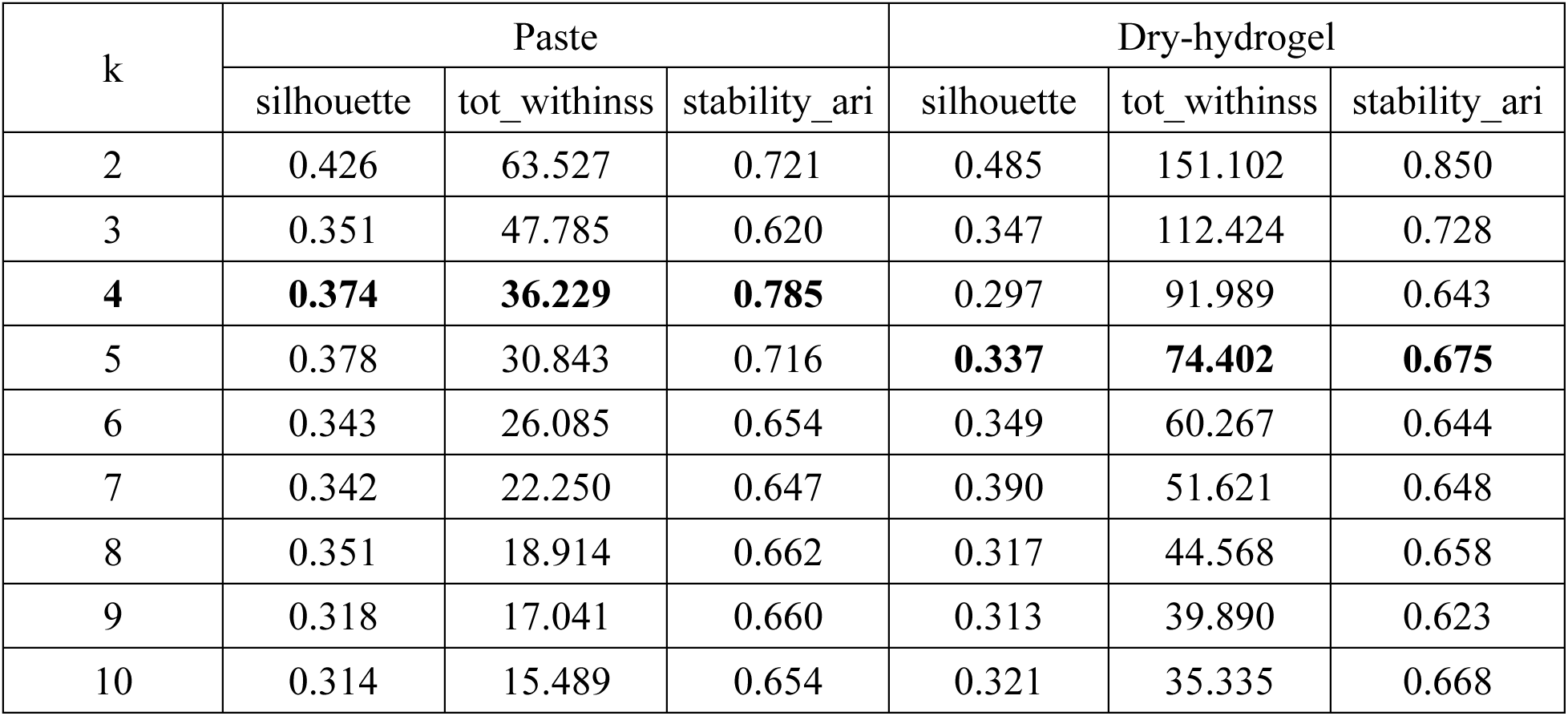
Cluster quality metrics for k-means clustering of physiological endpoints. For each candidate number of clusters (k = 2 to 10), we report the average silhouette width (higher values indicate better separation), the total within-cluster sum of squares (lower values indicate greater compactness), and the stability of the cluster solution assessed by the adjusted Rand index (ARI) after bootstrap resampling (values closer to 1 indicate higher reproducibility). The analysis was performed on the three-dimensional effect-size space (SF–VIV, SF–GC, SF-time). Based on these metrics, the optimal number of clusters (bold) for the paste and dry-hydrogel–fed cohorts

**Table S5.** Cluster-specific network metrics for paste-fed and dry-fed mice. For each cluster identified in Fig. 3, the table reports the number of nodes (endpoints), number of edges (correlations |r| ≥ 0.9), density, average and maximum degree, average and median within-cluster correlation, and a cohesion score (density × avg correlation). Higher cohesion indicates more tightly integrated functional modules

| Diet | Paste |  |  |  | Dry |  |  |  |  |
| --- | --- | --- | --- | --- | --- | --- | --- | --- | --- |
| Cluster | 1 | 2 | 3 | 4 | 1 | 2 | 3 | 4 | 5 |
| N_nodes | 16 | 13 | 13 | 31 | 24 | 3 | 25 | 13 | 8 |
| N_edges | 80 | 42 | 30 | 175 | 97 | 3 | 206 | 73 | 11 |
| Density | 0.667 | 0.538 | 0.385 | 0.376 | 0.351 | 1 | 0.687 | 0.936 | 0.393 |
| Avg degree | 10 | 6.46 | 4.62 | 11.29 | 8.08 | 2 | 16.48 | 11.23 | 2.75 |
| Max degree | 12 | 9 | 6 | 17 | 13 | 2 | 22 | 12 | 4 |
| Avg correlation | 0.901 | 0.837 | 0.754 | 0.664 | 0.708 | 0.998 | 0.897 | 0.97 | 0.693 |
| Median correlation | 0.943 | 0.91 | 0.809 | 0.808 | 0.802 | 0.997 | 0.958 | 0.984 | 0.637 |
| Cohesion score | 0.625 | 0.497 | 0.355 | 0.364 | 0.337 | 0.667 | 0.659 | 0.864 | 0.344 |

**Table S6.** Cluster architecture of spaceflight effects in the Nrf2 modulation experiment. Endpoints were clustered in a three-dimensional space defined by: d_(SF–VIV_С57)_ — the effect of spaceflight in untreated C57BL/6 mice (baseline response); d_(C57–RTA flight)_ — the effect size induced by pharmacological Nrf2 activation (RTA-408) relative to C57BL/6 in spaceflight; d_(C57–Nrf2KO flight)_ — the change in effect size in Nrf2-knockout mice relative to C57BL/6 relative to C57BL/6 in spaceflight. Positive *d* values indicate that the treatment or genotype attenuated the flight effect (pushed it toward control values); negative values indicate exacerbation. For each cluster, the table lists the composition, bootstrap stability (mean ± SD), and the mean values of the three clustering coordinates. This approach reveals how graded Nrf2 activity (disrupted, baseline, enhanced) selectively tunes distinct physiological modules without fundamentally rewiring the overall response architecture.

| Cluster | Composition | Bootstrep<br>stability,<br>m±sd | Mean Cohen's d |  |  |
| --- | --- | --- | --- | --- | --- |
|  |  |  | d<sub>(SF-VIV_C57)</sub> | d<sub>(C57-RTA flight)</sub> | d<sub>(C57-Nrf2KO flight)</sub> |
| Paste |  |  |  |  |  |
| 1<br>Stress-hematologic | Adrenal, ALT, AST, Ca/P, CK, ColonL, Epidyd, Fat, HCT, HGB, IntestL, LDH, MCH, MCHC, MCV, Mon#, Mon%, Neu#, Neu%, NLR, RBC, WBC | 89 ± 10 | 0.85 | -0.27 | -0.08 |
| 2<br>Muscle-core | Bulb, Gastrocnemius, Plantaris, Quad, RectusFem, Soleus, Tibialis | 97 ± 0 | -3.01 | -2.35 | -1.26 |
| 3<br>Visceral-organs | A/G, Eos#, Eos%, Heart, Intermedialis, Kidney, Lateralis, Liver, LPS, Pancreas, RDW-SD, Rventr, Septum, Spleen, Testes, TG | 95 ± 2 | -0.75 | -0.76 | -0.57 |
| 4<br>Visceral-metabolic | ALB, ALP, AMY, ColonM, Crea, Gaster, GLOB, GLU, MPV, PDW, TC, TP | 98 ± 1 | 1.09 | 1.25 | 1.27 |
| 5<br>Resilient-visceral | AccessGl, Bladder, Brain, BUN, BUN/CREA, Ca, IntestM, Lung, Lventr, Lym#, Lym%, Medialis, PHOS, PLT, Salivary, TB, tCO2, Thymus | 98 ± 1 | -0.52 | 0.10 | 0.35 |

**Table S7.** Cluster quality metrics for k-means clustering in the Nrf2 modulation experiment. Silhouette width and total within-cluster sum of squares (tot_withinss) for candidate numbers of clusters (k = 2 to 10). Higher silhouette values indicate better separation; lower tot_withinss indicates greater compactness.

| k | silhouette | tot_withinss |
| --- | --- | --- |
| 2 | 0.516 | 155.6 |
| 3 | 0.398 | 103.6 |
| 4 | 0.333 | 78.6 |
| **5** | **0.308** | **66.8** |
| 6 | 0.315 | 58.0 |
| 7 | 0.303 | 50.7 |
| 8 | 0.287 | 43.6 |
| 9 | 0.315 | 38.7 |
| 10 | 0.314 | 34.7 |

**Data S1. Term enrichment analysis (GSEA-like) of functional terms for paste-fed and dry-fed mice across three spaceflight contrasts**

For each diet, enrichment was calculated separately for three orthogonal contrasts: flight vs ground control (d SF–GC), flight vs vivarium control (d SF–VIV), and the within-flight recovery slope (dSF–time). For each functional term, the table reports the number of endpoints contributing to the term (N), the enrichment score (ES), the normalized enrichment score (NES), the nominal p-value (P_RAW_), and the false discovery rate (P_FDR_). Positive NES indicates enrichment toward higher values in the flight group (or positive recovery slope); negative NES indicates enrichment toward lower values. Terms with |NES| > 1 and P_FDR_ < 0.2 were considered biologically interpretable. The analysis reveals diet-dependent remodeling of functional architecture, with muscle-related terms consistently negative across contrasts and immune-inflammatory terms switching from negative (paste) to positive (dry) enrichment

**Data S2. Body weight, organ weights, hematology, biochemistry and statistical analysis**

Excel workbook with three worksheets containing raw measurements for all animals and groups, body-mass- and adiposity-adjusted organ weights, and results of all statistical analyses (linear models, ANOVA, pairwise contrasts and Cohen’s d).

**Data S3. Body weight, organ weights, hematology, biochemistry and statistical analysis for Nrf2 experiment**

Excel workbook with three worksheets containing raw measurements for all Nrf2 treatment groups (Nrf2-KO, C57BL/6, RTA-408), body-mass- and adiposity-adjusted organ weights, and results of all statistical analyses including linear models, ANOVA, pairwise contrasts and Cohen’s *d* for flight vs control comparisons across Nrf2 activity states.

